# The SEEL Motif and Members of the MYB-related REVEILLE Transcription Factor Family are Important for the Expression of *LORELEI* in the Synergid Cells of the Arabidopsis Female Gametophyte

**DOI:** 10.1101/2021.08.17.456723

**Authors:** Jennifer A. Noble, Alex Seddon, Sahra Uygun, Ashley Bright, Steven E. Smith, Shin-han Shiu, Ravishankar Palanivelu

## Abstract

Synergid cells in the micropylar end of the female gametophyte are required for critical cell-cell signaling interactions between the pollen tube and the ovule that precede double fertilization and seed formation in flowering plants. *LORELEI* (*LRE*) encodes a GPI-anchored protein that is expressed primarily in the synergid cells, and together with FERONIA, a receptor-like kinase, it controls pollen tube reception by the receptive synergid cell. Still, how *LRE* expression is controlled in synergid cells remains poorly characterized. We identified candidate *cis*-regulatory elements enriched in *LRE* and other synergid cell-expressed genes. One of the candidate motifs (‘*TAATATCT*’) in the *LRE* promoter was an uncharacterized variant of the Evening Element motif that we named as the Short Evening Element-like (SEEL) motif. Deletion or point mutations in the SEEL motif of the *LRE* promoter resulted in decreased reporter expression in synergid cells, demonstrating that the SEEL motif is important for expression of *LRE* in synergid cells. Additionally, we found that *LRE* expression is decreased in the loss of function mutants of REVEILLE (RVE) transcription factors, which are clock genes known to bind the SEEL and other closely related motifs. We propose that RVE transcription factors regulate *LRE* expression in synergid cells by binding to the SEEL motif in the *LRE* promoter. Identification of a *cis*-regulatory element and transcription factors involved in the expression of *LRE* will serve as a foundation to characterize the gene regulatory networks in synergid cells and investigate the potential connection between circadian rhythm and fertilization.

**One sentence summary:** A newly identified SEEL motif in the promoter of *LORELEI* and at least three members of the REVEILLE transcription factor family are important for *LORELEI* expression in synergid cells of the Arabidopsis female gametophyte.

## Introduction

Seeds are the principle propagules of angiosperms and gymnosperms and are important sources of food, fiber, feed, industrial products, oils, and biofuels. Seeds form when plants reproduce sexually and depend on interactions between the haploid male and female gametophytes (Johnson et al. 2019). In *Arabidopsis thaliana* (hereafter, Arabidopsis), the mature male gametophyte consists of two gametic cells (two sperm cells) and one accessory cell in the pollen tube (PT). The mature female gametophyte (FG) consists of two gametic cells (one egg cell and one central cell) and five accessory cells (three antipodal cells and two synergid cells) (Yadegari and Drews 2004).

Synergid cells control the final steps of the sexual plant reproduction by attracting the PT into the ovule (PT attraction), receiving the PT and inducing its lysis (PT reception) and enabling discharge of sperm cells (Huck et al. 2003; Sandaklie-Nikolova et al. 2007; Okuda et al. 2009; Amien et al. 2010; Hamamura et al. 2011; Leydon et al. 2015), which will then fuse with the egg and central cells to complete double fertilization. The fertilized ovule subsequently develops into a seed. Genes and the molecular mechanisms by which synergid cell-expressed genes control these indispensable events in plant reproduction are beginning to be understood through forward and reverse genetic analysis.

Mutants in synergid cell-expressed genes show aberrant PT attraction and PT reception defects, validating that synergid cells are critical for these final steps in plant reproduction (Huck et al. 2003; Rotman et al. 2003; Kasahara et al. 2005; Escobar-Restrepo et al. 2007; Capron et al. 2008; Kessler et al. 2010; Tsukamoto et al. 2010; Takeuchi and Higashiyama 2012; Duan et al. 2014; Hou et al. 2016; Liu et al. 2016; Zhong and Qu 2019). Profiling the synergid cellspecific transcriptomes in Arabidopsis and rice have revealed thousands of synergid cell-expressed and synergid cell-enriched genes, including transcription factors (TFs) (Wuest et al. 2010; Ohnishi et al. 2011). However, only a handful of TFs have been analyzed further and shown to control the expression of genes in synergid cells (Kasahara et al. 2005; Jones-Rhoades et al. 2007; Punwani et al. 2007; Punwani et al. 2008; Wang et al. 2010; Kirioukhova-Johnston et al. 2019), highlighting the need to establish a synergid cell gene regulatory network (GRN) by identifying additional TFs and binding motifs in their regulatory targets. This GRN will then lay the foundation to better understand the role of synergid cell-expressed genes in synergid functions, a critical need in our understanding of plant reproduction. In this study, we investigated *cis*-regulatory elements (CREs) and TFs that control the expression of *LORELEI* (*LRE*), a predominantly synergid cell-expressed gene in Arabidopsis.

LRE is a putative GPI-anchored membrane protein that functions prior to and after PT arrival (Capron et al. 2008; Tsukamoto et al. 2010; Li et al. 2015; Liu et al. 2016). Before PT arrival, LRE chaperones FERONIA (FER) through the synergid cell endomembrane system to the filiform apparatus (FA), a highly invaginated, membrane-rich region shared by both synergid cells, and play a pivotal role in PT-synergid cell interactions (Li et al. 2015). FER is a member of the *Catharanthus roseus* receptor-like kinase 1-like (CrRLK1L) subfamily in Arabidopsis and is expressed in synergid cells (Escobar-Restrepo et al. 2007). After localization in the FA, FER and LRE are important for reactive oxygen species (ROS) accumulation in the FA (Duan et al. 2014). Upon PT arrival, LRE and FER together trigger a distinct change in calcium signaling in the receptive synergid cell and induce PT reception (Ngo et al. 2014; Li et al. 2015; Liu et al. 2016).

*LRE* has three homologs in Arabidopsis, *LORELEI-LIKE GPI-ANCHORED PROTEIN 1, 2*, and *3* (*LLG1*, *LLG2*, and *LLG3*) (Tsukamoto et al. 2010). LLG1 is the most closely related to LRE (Tsukamoto et al. 2010; Noble et al. 2020) and functions with FER in vegetative tissues to regulate growth and development (Li et al. 2015). *LLG1* is expressed in many tissues throughout plant development, while *LLG2* and *LLG3* are most strongly expressed in the male gametophyte (Tsukamoto et al. 2010; Feng et al. 2019; Ge et al. 2019; Xiao et al. 2019). Unlike its homologs, *LRE* expression is mostly limited to the female gametophyte, where it is strongly expressed in synergid cells and weakly expressed in egg and central cells before fertilization (Wang et al. 2017). In fertilized seeds, *LRE* expression is imprinted, as only the matrigenic *LRE* allele is expressed for a short duration in the zygote and proliferating endosperm (Wang et al. 2017).

In this study, we used a bioinformatics approach to identify candidate CREs that control *LRE* expression in synergid cells before fertilization. By deleting or altering the sequence of a novel Short Evening Element-like (SEEL) motif, ‘*TAATATCT*’, in the *LRE* promoter of a *pLRE::GFP* transcriptional reporter fusion construct, we demonstrated that the SEEL motif was important for controlling *LRE* expression in synergid cells. In Arabidopsis, members of the REVEILLE (RVE) TF family are known to bind the SEEL motif and other variants of the Evening Element (EE) (Alabadi et al. 2001; Gong et al. 2008; Rawat et al. 2009; Rawat et al. 2011; Hsu et al. 2013; Jiang et al. 2016; O’Malley et al. 2016). Consistent with this, we found that GFP expression in synergid cells of plants carrying the *pLRE::GFP* is decreased in loss of function *rve1, rve5*, and *rve6* mutants. Findings from this work will facilitate characterization of the GRN in synergid cells and help identify novel proteins that function with LRE in PT reception, as expression of genes functioning in a pathway tends to be coregulated.

## Results

### The *LORELEI* promoter contains an eight base pair sequence that is a variant of the ‘Evening Element’ motif

To find putative CREs in the *LRE* promoter that control *LRE* expression in synergid cells, we used two approaches. In both approaches, we used an *in vitro* Transcription Factor Binding Motif (TFBM) dataset (Weirauch et al. 2014) and computationally-derived motif data (see Materials & Methods). *Cis*-regulatory sequences bound by TFs are preferentially located in the proximal region of the promoter, which is typically about 500 bp upstream of the transcription and/or translation start site (Zou et al. 2011; Franco-Zorrilla et al. 2014). Therefore, we focused on TFBMs in the proximal region of the *LRE* promoter. In the first approach, we identified TFBM sites that are present only in the *LRE* promoter but not in the promoters of other three members of the Arabidopsis *LLG* family (*LLG1, LLG2*, and *LLG3*), as only *LRE* is primarily expressed in the synergid cells of the Arabidopsis female gametophyte (Tsukamoto et al. 2010; Wang et al. 2017; Feng et al. 2019; Xiao et al. 2019; Noble et al. 2020). Using this approach, we identified three TFBMs in the proximal region *LRE* promoter but not in the proximal region of the promoter of its homologs: ‘*NATNATTNN*’, ‘*NNNTAWATTATNN*‘, and ‘*WAATATCT*’ (Supplementary Dataset 1). In the second approach, which was based on analyzing co-expression of synergid cell-expressed genes, we combined a heterogeneous expression dataset (see Materials and Methods) and used 5,446 synergid cell-expressed genes (Wuest et al. 2010) to generate 145 co-expression clusters (Supplementary Dataset 2). We found 49 putative CREs to be enriched in the promoters of synergid cell-expressed genes within these clusters (Supplementary Datasets 2 and 3). Two of these 49 enriched co-expression-derived motifs were present in the proximal region of the *LRE* promoter: ‘*NNNAAMGN*’ and ‘*WAATATCT*’ (Supplementary Datasets 2 and 3). As both approaches identified the ‘*WAATATCT*’ motif (where W is either A or T), we analyzed this motif in greater detail.

One of the two possible versions of the ‘*WAATATCT*’ motif, ‘*AAATATCT*’, was present in the promoters of approximately 25% of synergid cell-expressed genes within these co-expression clusters (Supplementary Dataset 4). The ‘*AAATATCT*’ motif is commonly known as the ‘Short Evening Element’ (SEE) (Hsu et al. 2013), which is a shorter variant of the Evening Element (EE) motif: ‘*AAAATATCT*’ (Harmer et al. 2000) (Table 1). The other possible version of the ‘*WAATATCT*’ motif, ‘*TAATATCT*’, was identified once in the *LRE* promoter, 229 bp upstream of the *LRE* translation start site. As this motif has not been reported previously, we named ‘*TAATATCT*’ as the ‘Short Evening Element-Like’ (SEEL) motif (Table 1).

**Table 1.**
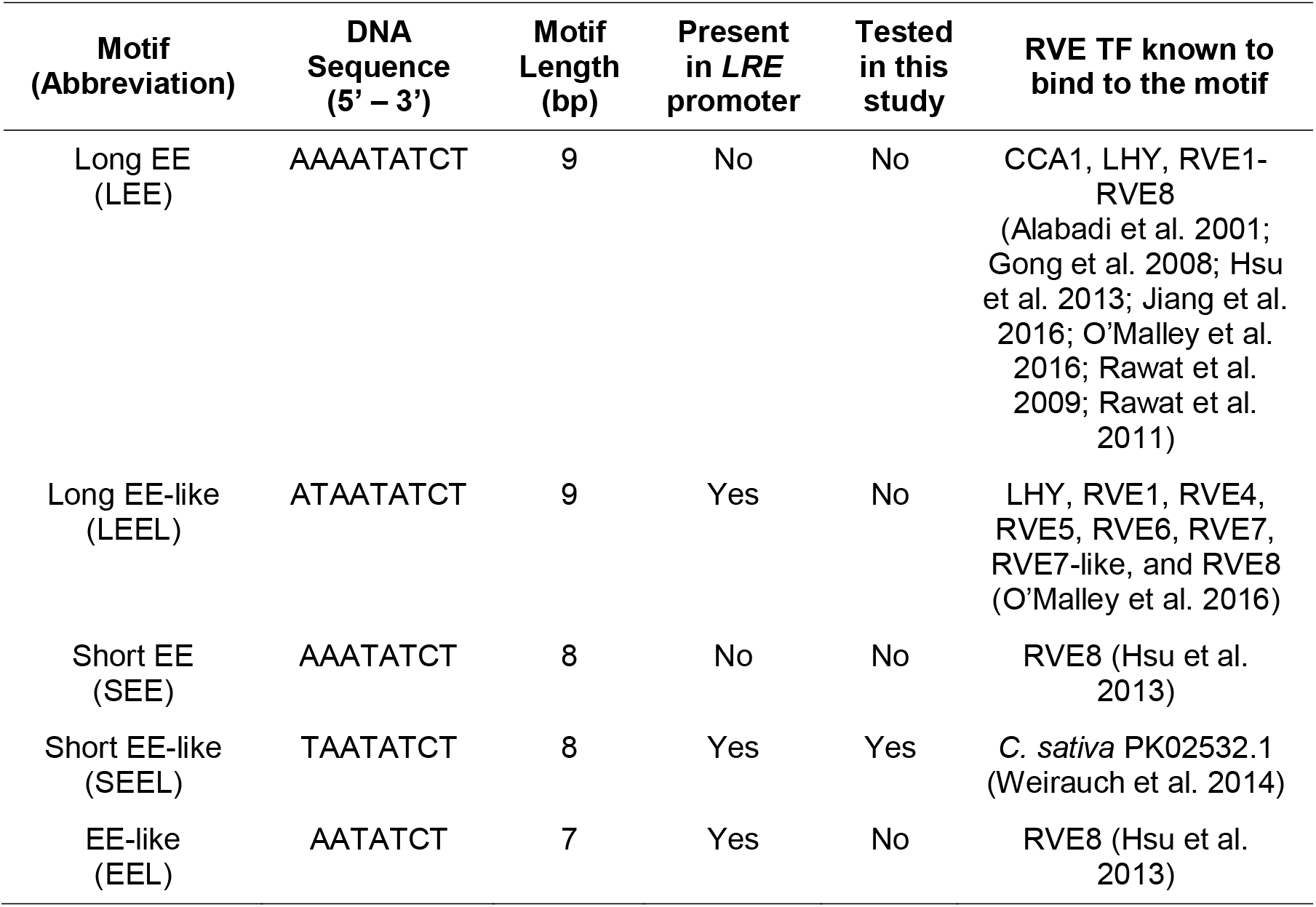
The Short Evening-Element Like (SEEL) motif identified using the CIS-BP database is related to the previously characterized EE motif

| Motif<br>(Abbreviation) | DNA<br>Sequence<br>(5' – 3') | Motif<br>Length<br>(bp) | Present<br>in *LRE*<br>promoter | Tested<br>in this<br>study | RVE TF known to<br>bind to the motif |
| --- | --- | --- | --- | --- | --- |
| Long EE<br>(LEE) | AAAATATCT | 9 | No | No | CCA1, LHY, RVE1-RVE8<br>(Alabadi et al. 2001;<br>Gong et al. 2008; Hsu<br>et al. 2013; Jiang et al.<br>2016; O'Malley et al.<br>2016; Rawat et al.<br>2009; Rawat et al.<br>2011) |
| Long EE-like<br>(LEEL) | ATAATATCT | 9 | Yes | No | LHY, RVE1, RVE4,<br>RVE5, RVE6, RVE7,<br>RVE7-like, and RVE8<br>(O'Malley et al. 2016) |
| Short EE<br>(SEE) | AAATATCT | 8 | No | No | RVE8 (Hsu et al.<br>2013) |
| Short EE-like<br>(SEEL) | TAATATCT | 8 | Yes | Yes | *C. sativa* PK02532.1<br>(Weirauch et al. 2014) |
| EE-like<br>(EEL) | AATATCT | 7 | Yes | No | RVE8 (Hsu et al.<br>2013) |

We next examined if the SEEL motif is conserved in the promoters of putative orthologs of *LRE* by analyzing promoter sequences for *LRE* orthologs from eleven species of Brassicaceae (see Materials and Methods). Indeed, the SEEL motif was present in the proximal promoter region of putative *LRE* ortholog in *Sisymbrium irio* and the related SEE motif was present in the proximal promoter region of the putative *LRE* orthologs in *Arabidopsis lyrata* and *Camelina sativa* (Table 2). Conservation of the SEEL motif in the promoters of these *LRE* orthologs raises the possibility that this motif may have a role in the expression of *LRE* in synergid cells.

**Table 2.** The SEEL motif and other EE variants are conserved in the promoters of putative orthologs of *LRE* in Brassicaceae

| Motif | Sequence Name | Star<sup>#</sup> | Stop<sup>#</sup> | Strand | Score | *p* value | *q* value | Promoter region |
| --- | --- | --- | --- | --- | --- | --- | --- | --- |
| SEEL | *Arabidopsis thaliana pLRE* | 722 | 730 | minus | 13.21 | 5.30 e-05 | 0.12 | Proximal |
| SEEL | *Brassica rapa pLRE* | 212 | 220 | plus | 13.46 | 2.39 e-05 | 0.12 | Distal |
| SEEL | *Leavenworthia alabamica pLRE* | 24 | 32 | plus | 13.21 | 5.30 e-05 | 0.12 | Distal |
| SEEL | *Leavenworthia alabamica pLRE* | 31 | 39 | plus | 13.21 | 5.30 e-05 | 0.12 | Distal |
| SEEL | *Sisymbrium irio pLRE* | 896 | 904 | plus | 13.46 | 2.39 e-05 | 0.12 | Proximal |
| SEE | *Arabidopsis lyrata pLRE* | 801 | 809 | minus | 13.21 | 5.30 e-05 | 0.12 | Proximal |
| SEE | *Brassica rapa pLRE* | 93 | 101 | minus | 13.21 | 5.30 e-05 | 0.12 | Distal |
| SEE | *Camelina sativa pLRE-1* | 242 | 250 | minus | 13.21 | 5.30 e-05 | 0.12 | Distal |
| SEE | *Camelina sativa pLRE-1* | 738 | 746 | minus | 13.21 | 5.30 e-05 | 0.12 | Proximal |
| EEL | *Aethionema arabicum pLRE* | 244 | 252 | minus | 9.20 | 9.66 e-05 | 0.19 | Distal |
| EEL | *Brassica rapa pLRE* | 452 | 460 | minus | 9.20 | 9.66 e-05 | 0.19 | Distal |

<sup>#</sup>Start and stop refer to the nucleotide position in a promoter of a gene, which is defined as 1000bp upstream of the translation start site of that gene. Positions greater than 500 indicate presence in the proximal promoter region, beyond that they were in the distal promoter region. Motifs were mapped using Find Individual Motif Occurrences (FIMO Version 5.0.5).
Only the *p* values that are statistically significant and less than 1e-4 are reported in this table. *q* values (false discovery rate at which a motif occurrence is significant) are reported by FIMO Version 5.0.5 only if the *p* values were deemed statistically significant. They were determined by the Benjamini and Hochberg method (Benjamini and Hochberg 1995).

### The 8bp SEEL motif in the *LRE* promoter is important for *LRE* expression in synergid cells

To test if the SEEL motif is important for *LRE* expression in synergid cells, we mutated the SEEL motif in the *pLRE::GFP* reporter construct (Wang et al. 2017). In the first mutant construct, we deleted the SEEL motif in the *LRE* promoter (*pLREΔSEEL::GFP*) (Fig. 1A). In two other mutant constructs, we altered the SEEL motif sequence such that either the sequence and the GC composition of the motif were altered (*pLRE-m1-SEEL::GFP*) or only the sequence of the motif was changed (*pLRE-m2SEEL::GFP*) (Fig. 1A). The three mutant constructs were transformed into wild-type Arabidopsis plants and the GFP expression in synergid cells was compared to that in transformants carrying the unmutated *pLRE::GFP* construct.

**Fig. 1.**
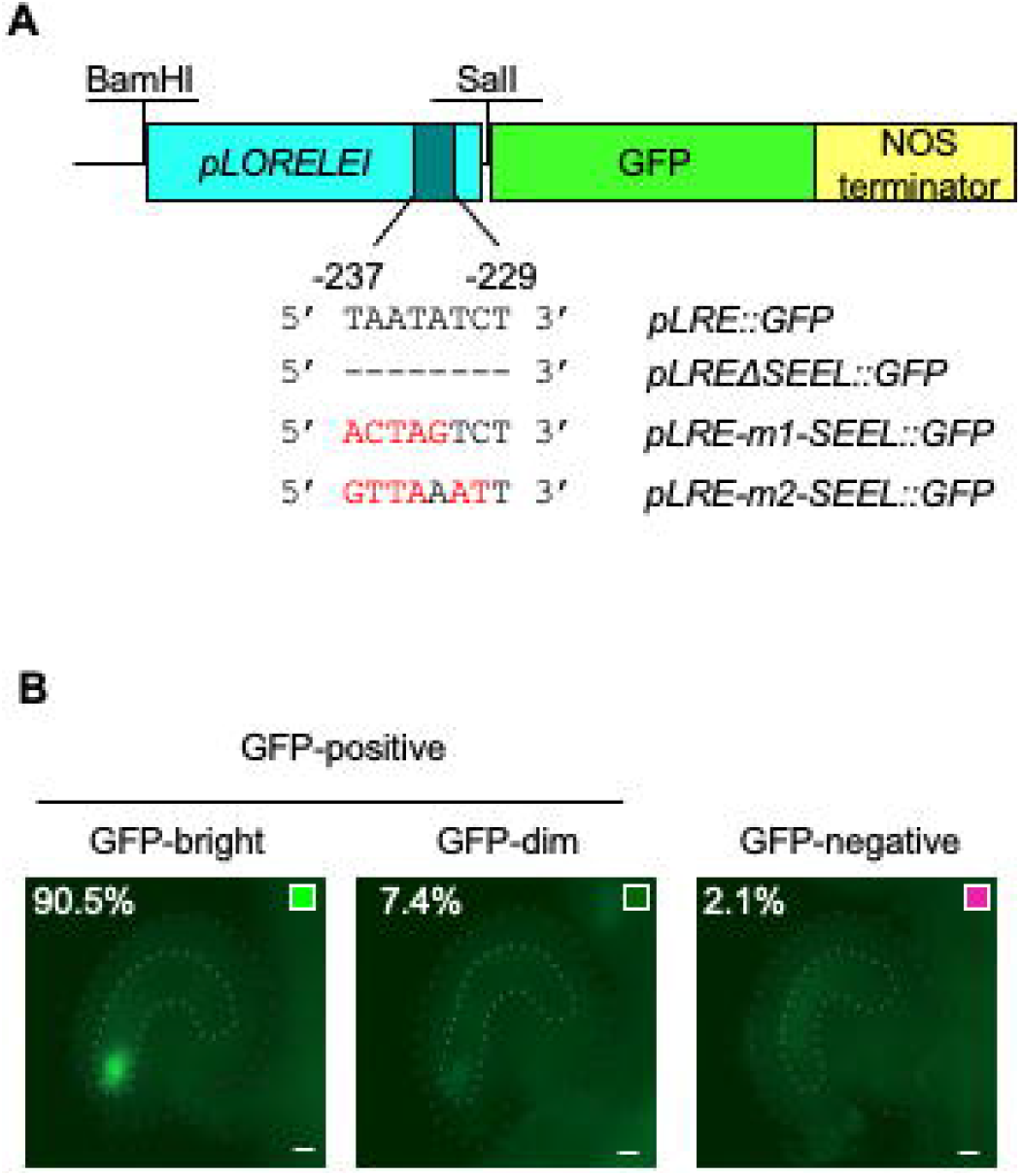
Constructs and synergid cell expression assay used in this study to examine the importance of the SEEL motif in the expression of LORELEI in synergid cells. A. Diagram of motif mutation and deletion constructs. The SEEL motif is present −237bp to −229 bp upstream of the translation start site of *LORELEI*. B. *pLRE::GFP* is expressed in the synergid cells of the female gametophyte located within an ovule and intensity of GFP was categorized as GFP–negative (pink, ovules with no GFP), GFP-dim (dark green, ovules with low GFP), or GFP-bright (bright green, ovules with high GFP). Images shown here were captured using identical camera settings. Scale bar: 10μm

To gain initial insights into the GFP expression from the reporter transgenes, ovules of T1 lines were scored as either GFP-positive or GFP-negative. Two sets of GFP expression data in single-locus insertion, hemizygous T1 lines showed that either deleting or mutating the SEEL motif in the *pLRE::GFP* construct reduced the GFP reporter expression in the synergid cells (Fig. 2). First, there was a decrease in the number of single-locus insertion, hemizygous mutant SEEL motif T1 lines with > 35% GFP-positive ovules in a pistil, a rate of GFP expression detected in single-locus insertion, hemizygous unmutated *pLRE::GFP* T1 lines (Fig. 2A; Table 3). Second, compared to T1 lines carrying the *pLRE::GFP* transgene, the T1 lines carrying mutant SEEL motif transgenes showed a decrease in the number of GFP-positive ovules. The median percentage of GFP-positive ovules in pistils of single-locus insertion T1 lines carrying the *pLREΔSEEL::GFP* transgene had a significantly lower proportion of GFP-positive ovules compared to *pLRE::GFP* T1 lines (median = 0%; Mann-Whitney *U*-test, *p* = 7.3e-4; Fig. 2B; Table 3). Similarly, the median percentage of GFP-positive ovules among single insertion T1 lines carrying the *pLRE-m1-SEEL::GFP* transgene was 0%, which was significantly lower than that for the *pLRE::GFP* control construct (*p* = 2.4e-3; Fig. 2C; Table 3). The single-locus insertion T1 lines carrying the *pLRE-m2-SEEL::GFP* transgene also had reduced GFP expression in ovules but this reduction was less severe compared to the other two mutant constructs (median = 6%; *p* = 5.4e-2; Fig. 2D; Table 3).

**Fig. 2.**
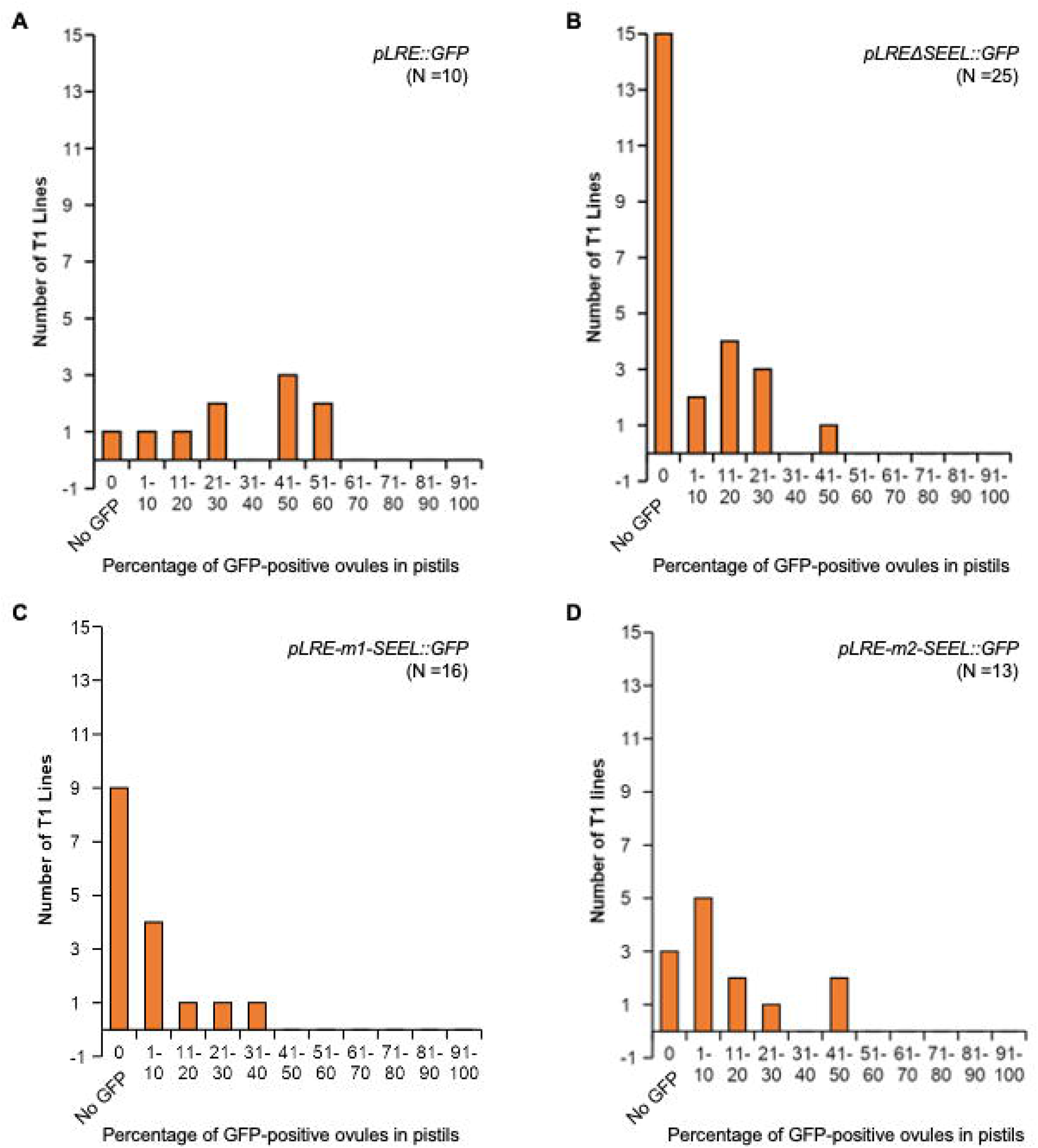
Majority of single-locus insertion T1 lines carrying the SEEL motif deletion or alteration showed a reduction in the percentage of GFP-positive ovules In T1 plants carrying *pLRE::GFP* (A) *pLREΔSEEL::GFP* (B), *pLRE-m1-SEEL::GFP* (C), and *pLRE-m2-SEEL::GFP* (D) constructs, GFP was scored in mature ovules, 24 hours after emasculation. Single-locus insertion lines were identified by segregation ratios of hygromycin resistance in T2 seedlings raised from selfed seeds of T1 lines. N equals the total number of T1 single insertion lines scored for each construct. Each T1 plant represents a single transgenic line, in which ovules from 2-3 emasculated pistils were scored.

**Table 3.** Single-locus insertion T1 lines with changes to the SEEL motif in the promoter of the *pLRE::GFP* construct show decreased numbers of GFP-positive ovules compared to control *pLRE::GFP* T1 lines

| Genotype | Fraction of T1 lines with GFP-positive ovules | Fraction of T1 lines with >40% GFP-positive ovules in a pistil | Median % GFP-positive ovules in pistils of T1 lines |
| --- | --- | --- | --- |
| *pLRE::GFP* | 9/10 | 5/9 | 35.7% |
| *pLREΔSEEL::GFP* | 10/25 | 1/10 | 0%** |
| *pLRE-m1-SEEL::GFP* | 7/16 | 0/7 | 0%** |
| *pLRE-m2-SEEL::GFP* | 10/13 | 2/10 | 6%** |

To test if decrease in GFP expression in synergid cells of ovules was also observed in subsequent generations of lines carrying the mutant transgenes, we selected four single-locus insertion T1 lines for each construct (three randomly selected T1 lines and the fourth was a T1 line with the highest percentage of GFP-positive ovules in Fig. 2). In the subsequent generations of each of these four lines, we identified plants homozygous for the transgene, sequence verified the transgene, and scored either GFP-negative or GFP-positive ovules and the latter was classified further as GFP-bright or GFP-dim (Fig. 1B). As seen in the T1 generation, the T2/T3 generation *pLRE::GFP* lines also had higher percentages of GFP-positive ovules (ranging from 92.5% to 98.0%, after combining GFP-bright and GFP-dim ovules) (Fig. 3). Among the four *pLREΔSEEL::GFP* lines scored, the mean percentages of ovules with GFP-negative, GFP-dim, and GFP-bright were 75.4%, 6.8%, and 17.8%, respectively. The percentages of GFP-positive ovules in these lines were significantly lower compared to those in the *pLRE::GFP* control construct (*Chi*-square test, all control lines compared to all *pLREΔSEEL::GFP* lines, all multiple testing corrected *p* < 2.0e-6, Supplementary Table 1).

**Fig. 3.**
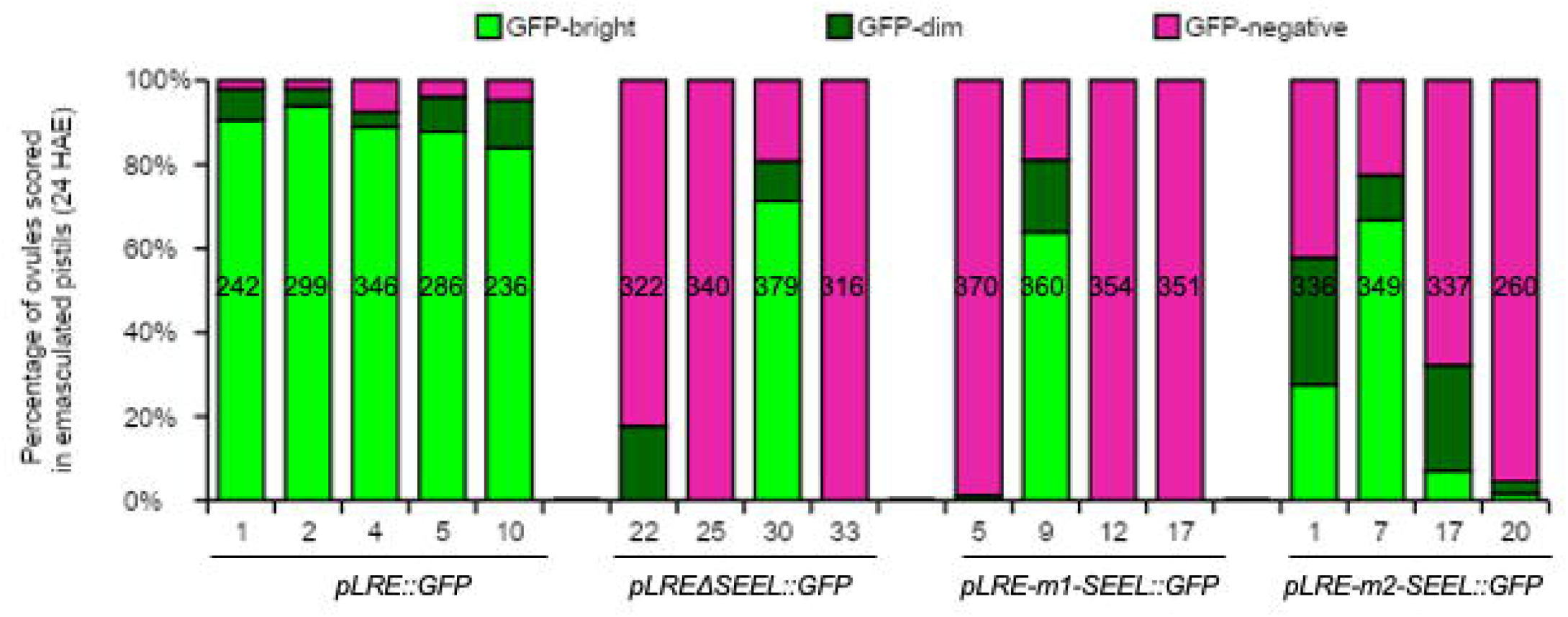
The SEEL motif is important for *LORELEI* expression in synergid cells. GFP was scored in mature ovules, 24 hours after emasculation in four homozygous single insertion lines in each of the three indicated mutant constructs. Line numbers are indicated in the X-axis. Five *pLRE::GFP* lines were scored, of which one is a previously published line (Wang et al. 2017). In each line, nine pistils from three homozygous plants (three pistils per plant) were scored and the total number of ovules scored for each line is indicated in the middle of each column. Statistical significance of decrease in GFP-positive ovules in each mutant line compared to that in each control line (*pLRE::*GFP) was evaluated using a *Chi*-square test and *p* values are reported in Supplementary Table 1.

The lines carrying other two SEEL motif mutant constructs also showed a noticeably decreased percentages of GFP-positive ovules and higher numbers of GFP-negative ovules compared to those in *pLRE::GFP* lines. In the four *pLRE-m1-SEEL::GFP* lines, an average of 79.4%, 4.6%, and 16.0% of ovules had GFP-negative, GFP-dim, and GFP-bright expression, respectively (Fig. 3). The percentages of GFP-positive ovules were significantly lower compared to the control line (all multiple control testing corrected *p* < 8.2e-8, Supplementary Table 1). All four *pLRE-m2-SEEL::GFP* lines also contained significantly lower percentages of GFP-positive ovules compared to those in *pLRE::GFP* transgenic lines (all multiple control testing corrected *p* < 2.4e-8, Supplementary Table 1). These results showed that the SEEL motif in the *LRE* promoter is important for *LRE* expression in synergid cells and validated bioinformatic predictions of a role for the SEEL motif in the expression of *LRE* in synergid cells.

### *LRE* expression is reduced in *rve* transcription factor mutant synergid cells

Regulation of gene expression involves binding of transcription factors to CREs (Zou et al. 2011; Franco-Zorrilla et al. 2014). To identify candidate TFs that may interact with the SEEL motif in the *LRE* promoter and control *LRE* expression, we first searched public databases for TFs that are known to bind the SEEL motif. Indeed, the Catalog of Inferred Sequence Binding Preferences (CIS-BP) database contained a MYB-related transcription factor in *Cannabis sativa* (PK02532.1) that was shown to bind the SEEL motif *in vitro* using its SHAQKYF-type of DNA-binding domain (DBD) (Weirauch et al. 2014). Predictions based on the homology of DBDs in the CIS-BP database (please see Materials and Methods) identified the Arabidopsis REVEILLE (RVE) TFs RVE3-6 and RVE8, which belong to the 11 member RVE TF family (Supplementary Fig. 1) that function as outputs and core components of the circadian clock and act as transcriptional activators or repressors (McClung, 2006; Hsu and Harmer, 2014). Indeed, the DBD in all 11 RVE TF family members shared a strong homology with the DBD of PK02532.1 (≥84.78% similarity and ≥58.67% identity) (Supplementary Fig. 2).

Analysis of published DNA-affinity purification sequencing (DAP-seq) data showed that RVE1, RVE4, RVE5, RVE6, RVE7, RVE7-like, RVE8, and closely related LHY bind to the SEEL motif and other variants listed in Table 1 (O’Malley et al. 2016). Searching the Plant Cistrome Database (http://neomorph.salk.edu/dap_web/pages/index.php) revealed that the Long Evening Element-like (LEEL) motif in the *LRE* promoter is a target for RVE1 and RVE5 binding *in vitro* (O’Malley et al. 2016). RVE1, RVE2, RVE3, RVE4, RVE7, RVE8, CCA1, and LHY are known to bind to the Long Evening Element (LEE) motif in protein binding microarrays and electrophoresis mobility shift assays (Alabadi et al. 2001; Gong et al. 2008; Rawat et al. 2009; Rawat et al. 2011). Finally, the EE-variants such as the SEE and EE-like (EEL) motifs were reported to be enriched in the promoters of differentially-expressed genes in *rve8* mutants (Hsu et al. 2013). Based on these observations, we hypothesized that the SEEL motif control of *LRE* expression in the synergid cells involve RVE TFs.

We tested if the *LRE* expression in synergid cells is affected in *rve* mutants, an outcome that could be expected if RVE TF plays a role in controlling *LRE* expression. To begin with, we used RT-qPCR to quantify endogenous *LRE* expression in *rve1, rve5*, and *rve6* mutant pistils for two reasons: first, of the 11 *RVE* genes in Arabidopsis, *RVE1, RVE5*, and *RVE6* genes are the three that were reported as ‘expressed’ in synergid cells in a microarray-based profiling of Arabidopsis synergid cell transcriptomes (Wuest et al. 2010). Second, RVE1, RVE5, and RVE6 are known to bind the LEEL motif (O’Malley et al. 2016), which encompasses the SEEL motif – the minimal novel motif that we identified to be important for *LRE* expression in synergid cells (Figs. 2 and 3). We obtained three alleles of *RVE1* (Supplementary Fig. 3A) and one mutant allele each of *RVE5* and *RVE6* (Supplementary Figs. 4A and 5A) and performed RT-qPCR experiments in mature unpollinated pistils (24 hours after emasculation, see Materials and Methods).

Compared to wild type, *RVE1* expression was nearly abolished in *rve1-1, rve1-2*, and *rve1-3* mutant pistils (Supplementary Fig. 3B; mean separation based on a mixed-model analysis of variance and a Tukey-Kramer test, *p < 1e-4*), *RVE5* and *RVE6* expression were significantly reduced in *rve5-1* mutant pistils (a decrease of 48.8% ± 0.16; Supplementary Fig. 4B; mean separation based on a Tukey-Kramer test, *p < 1e-4*) and *rve6-2* mutant pistils (a decrease of 86.1% ± 0.21; Supplementary Fig. 5B; mean separation based on a Tukey-Kramer test, *p < 1e-4*), respectively. These results confirmed that individual *rve* mutants are loss of function mutants in which corresponding *RVE* expression is significantly reduced and can therefore be useful in analyzing the role of RVE in controlling *LRE* expression.

We next performed RT-qPCR experiments to examine *LRE* expression using the same cDNAs from unpollinated pistils of *rve* mutants that were used to identify decreases in *RVE* expression. Compared to wild type, *LRE* expression decreased by 30.5% ± 0.11, 37.8% ± 0.11, and 22.8% ± 0.16 in *rve1-1, rve1-2*, and *rve1-3*, respectively (Supplementary Fig. 6). *LRE* expression was also reduced in *rve5-1* and *rve6-2* single mutant pistils, with decreases of 14.8% ± 0.22 and 43.1% ± 0.09, respectively, compared to wild type (Supplementary Fig. 6). Although these decreases of *LRE* expression in the different mutants were noticeable, the decreases were not statistically significant in any of the *rve* mutants compared to that in wild-type unpollinated pistils (Supplementary Fig. 6; mean separation based on Tukey-Kramer test; *p* = 5.0e-2). One reason for this could be that RT-qPCR analysis of unpollinated pistils is not sufficiently sensitive to detect changes in *LRE* expression in synergid cells (just two cells in an ovule where *LRE* is primarily expressed) (Wang et al. 2017), notwithstanding that we removed the stigma, style, and ovary walls before using the unpollinated pistils in the RT-qPCR experiments.

To address this caveat, we investigated if loss of function mutations in *RVE 1, RVE5*, and *RVE6* affected *LRE* expression specifically in synergid cells by crossing a well characterized single-locus insertion *pLRE::GFP* line (Line 1, Fig. 3 and (Wang et al. 2017)) to *rve5-1* and *rve6-2* single mutants, and two mutant alleles of *RVE1* (*rve1-1* and *rve1-2*) (Fig. 4). In each case, from the progeny of these crosses, we identified plants that were homozygous for both the *rve* mutation and the *pLRE::GFP* transgene and scored GFP expression in mature unpollinated pistils of these plants.

**Fig. 4.**
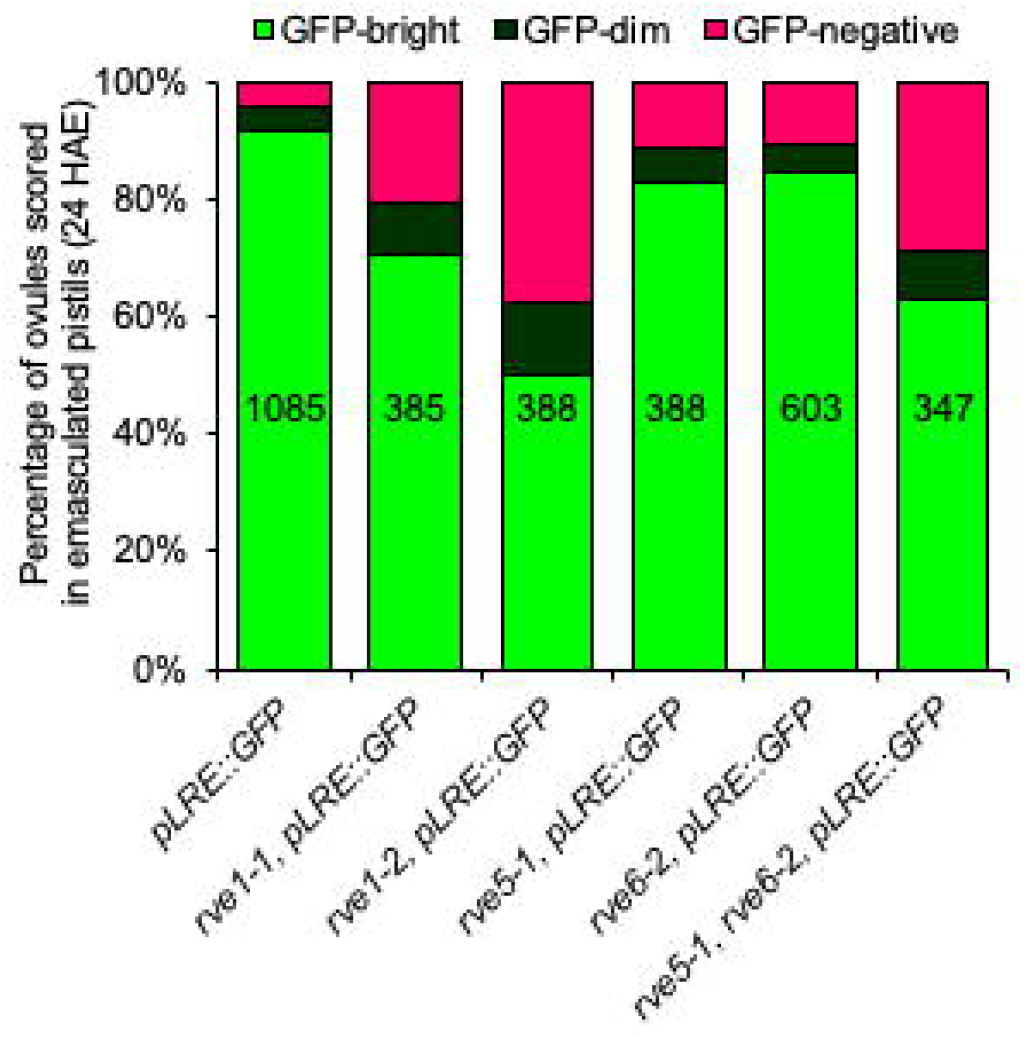
*pLRE::GFP* expression was decreased in unpollinated *rve* mutant pistils. In each of the indicated *rve* mutants, ovules from 9 to 25 mature unpollinated pistils (after removing ovary walls, stigma, style, and pedicel) were scored for GFP in the synergid cells of the female gametophyte, 24 hours after the pistils were emasculated. The total number of ovules scored for each mutant is shown in the middle of each column in the graph. Significance of difference in GFP expression between wild type and *rve* mutants was performed using a *Chi*-square test and the *p* values are reported in Supplementary Table 3.

Two sets of observations indicated that *rve* mutations affected *LRE* expression in synergid cells. First, in each of the *rve* mutants, the proportion of GFP-negative ovules was significantly increased compared to that in wild-type containing the *pLRE::GFP* transgene (Fig. 4; *Chi*-square test, *p > 1.0e-3;* Supplementary Table 3). Second, we observed changes in proportion of GFP-positive ovules in *rve* mutants expressing *pLRE::GFP*. In both *rve1-1 pLRE::GFP* and *rve1-2 pLRE::GFP* mutant pistils, we saw a significant decrease in percentages of GFP-bright ovules and a significant increase in percentages of GFP-dim ovules compared to that in wild-type plants carrying the *pLRE::GFP* transgene (Fig. 4, *Chi*-square test *p* values in Supplementary Table 3). In *rve5-1 pLRE::GFP* pistils and *rve6-2 pLRE::GFP* single mutant pistils, there were noticeable decreases and increases of percentages of GFP-bright ovules and GFP-dim ovules, respectively (Fig. 4); however, these decreases and increases were not significantly different compared to that in wild-type plants carrying the *pLRE::GFP* transgene (*Chi*-square test *p* values in Supplementary Table 3). RVE5 and RVE6 are most closely related to each other (Supplementary Fig. 1); therefore, we scored GFP-positive ovules in *rve5-1 rve6-2* double mutant pistils. Indeed, we found a significant decrease and increase in the proportion of GFP-bright ovules and GFP-dim ovules, respectively, compared to that in wild type, *rve5-1*, or *rve6-2* single mutant pistils expressing *pLRE::GFP* (Fig. 4; *Chi*-square test *p* values in Supplementary Table 3). In summary, increases in proportion of GFP-negative ovules, increases in proportion of GFP-dim ovules, and decreases in proportion of GFP-bright ovules in *rve* mutant ovules expressing *pLRE::GFP* led us to conclude that *LRE* expression is decreased in *rve1, rve5*, and *rve6* mutant synergid cells and suggested that *LRE* promoter is a likely target of *RVE1, RVE5*, and *RVE6* TFs.

## Discussion

### The SEEL motif is important but not essential for *LRE* expression in synergid cells

Our study determined that the SEEL motif in the *LRE* promoter is important for *LRE* expression in synergid cells. We observed a significant decrease in GFP-bright ovules in the *pLREΔSEEL::GFP* and *pLRE-m1-SEEL::GFP* lines when compared to the *pLRE-m2-SEEL::GFP* line, where the SEEL motif was mutated but the GC composition remained the same as the unmutated SEEL motif. Based on these observations, we propose that a combination of the sequence and GC composition of the motif is important for *LRE* expression. Our results also demonstrated that the SEEL motif was not essential because deleting or altering the SEEL motif significantly decreased the percentage of GFP-bright ovules but did not completely abolish *GFP* expression in synergid cells of all transgenic lines (Fig. 3). This raises the possibility that additional motifs in the *LRE* promoter control *LRE* expression in synergid cells. Consistent with this possibility, in our bioinformatic analysis we identified two other candidate motifs: ‘*NATNATTNN*’ and ‘*NNNAAMGN*’, which were present more than once in the *LRE* promoter (7 and 24 times, respectively). Additional promoter-deletion analysis will be required to test if these and additional CREs in the *LRE* promoter are important for *LRE* expression in synergid cells.

One other motif that is required and sufficient for synergid cell expression of genes is ‘*AACGT*’ which is part of the motif (GTAACNT) that interacts with the synergid cell-expressed MYB98 TF (Punwani et al. 2008). The MYB98 binding motif is present in some promoters of synergid cell-expressed cysteine-rich peptides (CRPs), involved in PT attraction (Punwani et al. 2008). Although SEEL motif is important for *LRE* expression in the synergid cells, it is unlikely to confer synergid cell-specific expression to a minimal promoter because many of the genes that carry this motif in their promoters are expressed in other cells besides synergid cells (Supplementary Dataset 4). Furthermore, *RVE* genes are expressed throughout plant development and also their protein products regulate expression of genes in vegetative tissues (Gray et al. 2017). Still, the SEEL motif is enriched in hundreds of synergid cell-expressed genes pointing to its importance in synergid cell expression.

### *FER* and *EVN*, genes that are expressed in synergid cells and function in PT reception, also contain a SEE(L) motif in their promoters

Besides *LRE*, the SEEL motif is present in many other genes that are expressed in synergid cells. We examined which of these genes are involved in PT reception to gain insights into the possibility that synergid cell-expressed genes that are co-regulated with *LRE* also function in the same pathway. For this, we checked if the SEEL motif is present in the promoters of genes known to be involved in PT reception - *FER*, *EVAN* (*EVN*), *ABSTINENCE BY MUTUAL CONSENT* (*AMC*), *TURAN* (*TUN*), and *EARLY NODULIN-LIKE14/15* (*EN14/15*) (Johnson et al. 2019). Of these genes, promoters of *AMC, TUN*, and *EN14/15* did not contain any SEEL motif or its variants. The SEE motif “AAATATCT” (Supplementary Table 2) is present once in the 5’UTR of *FER*. Additionally, the SEE motif is also conserved in the proximal region of promoters and/or 5’UTRs of putative *FER* orthologs in *Arabidopsis lyrata* and *Sisymbrium irio*, the two species which contain *LRE* orthologs that also have their SEE(L) motif conserved in their promoters (Supplementary Table 2). The SEE motif is also present once (>500bp upstream of the transcription start site) in the promoter of Arabidopsis *EVN*, which encodes a dolichol kinase that is required for biosynthesis of Dol-P and protein glycosylation; homozygous mutants are embryo lethal and heterozygous mutants show defects in PT reception in ovules and defects in pollen development (Kanehara et al. 2015; Lindner et al. 2015). We found that the SEEL motif was also present in the promoter of the putative *EVN* ortholog in *Sisymbrium irio*.

Analysis of the PT reception genes with and without the SEE(L) motif revealed an interesting pattern. The SEE(L) motif is absent in *AMC*, *TUN*, and *EN14/15* and there is no evidence linking them with *LRE* in a molecular function, even though these genes encode proteins that function in the developmental process of PT reception. However, *FER* and *EVN*, which contain a SEE(L) motif in their promoters, are linked with *LRE* in a molecular function during PT reception. FER and LRE bind with each other (Duan et al. 2014), LRE cochaperones FER to the FA (Li et al. 2015), both are co-receptors in the signaling pathway that controls reactive oxygen species production in the FA (Duan et al. 2014) and they both function together in calcium signaling in synergid cells upon PT arrival (Ngo et al. 2014). Furthermore, LRE is proposed to be a substrate of EVN (Lindner et al. 2015), as yeast *evn* mutant contains a dramatically reduced amount of GPI-anchored proteins (Heller et al. 1992). Based on this analysis, we propose that expression of *FER*, *EVN*, and *LRE* in synergid cells, which function together in PT reception, may be co-regulated.

### Partial reductions of *LRE* expression in *rve* mutants may be due to redundancy within the RVE TF family

Single mutants of *RVE1, RVE5*, and *RVE6* did not completely abolish *LRE* expression (Supplementary Fig. 6). One likely reason for this is that the *LRE* expression is controlled by additional TFs besides RVE. This is likely, as we know the SEEL motif is important but not essential for *LRE* expression in synergid cells. In addition to the SEEL motif, two other motifs in the *LRE* promoter were predicted: ‘*NATNATTNN*’ and ‘*NNNAAMGN*’. A Homeodomain Leucine Zip class TF (*At2g22430*) and a DOF Zinc finger TF (*At3g21270*), respectively, predicted to bind to these two motifs (Weirauch et al. 2014) are among the genes expressed in synergid cells (Wuest et al. 2010). Such TFs may provide additional control of *LRE* expression in the synergid cells.

Another reason why the single mutants that we investigated only show partial reductions in *LRE* expression could be due to the redundancy within the RVE TF family. All 11 members of the RVE TF family can bind to the EE motif and its variants (Table 1), and RVEs can be partially redundant in function (Mizoguchi et al. 2002; Hsu et al. 2013; Gray et al. 2017). Consistent with this possibility, a higher percentage of GFP-negative ovules were detected in the *rve5-1, rve6-2* double mutant compared to the single mutants (Fig. 4).

LHY and CCA1 are most closely related to each other and tend to act as repressors of genes with the EE motifs, while RVE8 and its most closely related homologs are typically transcriptional activators; however, their function as a repressor or activator is not mutually exclusive (Harmer and Kay, 2005; Rawat et al. 2009; Hsu et al. 2013). Based on decreased expression levels of *LRE* in *rve1, rve5*, and *rve6* mutants, we concluded that RVE1, 5, and 6 were likely transcriptional activators of *LRE* expression in synergid cells. Still, this hypothesis needs to be tested in more detail. Because RVEs can either serve as repressors or activators, higher order mutants can also have antagonistic interactions (Shalit-Kaneh et al. 2018) and thus prove difficult to analyze by having negligible net effect on *LRE* expression. Since *LRE* is expressed primarily in synergid cells, our study focused on *RVE1*, *RVE5*, and *RVE6*, which are known to be expressed in synergid cells (Wuest et al. 2010). However, this microarray-based expression data is not comprehensive; for example, *RVE7* and *RVE7-like* were not included in the microarrays used in the study by (Wuest et al. 2010) and additional experiments such as RNA-seq of synergid cells directly obtained using Laser Capture Microdissection (LCM) or by sorting of synergid cells tagged with transcriptional fusions of *RVE* TFs to reporter genes will be required.

### Synergid cell functions that are important for double fertilization may be circadian clock-dependent

In Arabidopsis, over one third of the genome is regulated by the circadian clock (Covington et al. 2008). Many of these genes contain the EE-motif or an EEvariant in their promoter, suggesting that the EE-motif is important for expression of circadian clock-regulated genes (Harmer et al. 2000). Members of the RVE TF family in Arabidopsis bind to the EE-motif in these genes and regulate gene expression in a circadian clock-dependent manner (Alabadi et al. 2001; Hsu et al. 2013). The function of RVEs is highly conserved throughout land plants and charophytes, suggesting they serve essential functions in regulating the expression of circadian clock-dependent genes (Linde et al. 2017). Since RVE roles are highly conserved, it is likely that the EE-motif is also highly conserved. Although further studies are required to test this possibility, the CIS-BP database already supports this proposal, as RVE-like TFs from other species can bind the SEEL motif (Weirauch et al. 2014).

The EE-motif and EE-variants have been identified in promoters of genes important for key processes, such as pathogen response and flowering (Lu et al. 2017; Inoue et al. 2018). It is likely that the EE-motif and its variants serve as master CREs to quickly induce or repress gene expression. Perhaps the SEEL motif is enriched in synergid cell-expressed genes to regulate or optimize synergid cell-expressed genes important for PT-ovule interactions.

We propose that PT-ovule interactions are regulated by the circadian clock, with a set of genes primed to be optimally expressed during the early evening phase of the clock. In this model, expression of synergid cell-expressed genes that are important for PT-ovule interactions peaks in the mid-afternoon and early-evening and render ovules at their performance maxima in the evening and receive PTs that started their journey on the stigma in the morning. Such a proposal is consistent with the findings that pollination peaks in the morning hours (van Doorn and Kamdee, 2014) and the elapsed time between the morning and evening is the time that it takes for PTs to reach ovules. By controlling the expression of genes involved in PT-ovule interactions using the SEEL motif in their promoters, perhaps optimal fertilization may be achieved during the evening phase of the circadian clock.

## Conclusions

In this study, we identified and validated a novel SEEL motif in the *LRE* promoter that is important for its expression in synergid cells and found that mutants of synergid cell-expressed RVEs decrease *LRE* expression in the synergid cells of unpollinated ovules. By identifying the SEEL motif in *LRE* and other synergid cell-expressed genes and implicating a role for RVE1 in *LRE* expression, our study will facilitate characterization of the GRN in synergid cells, critical cells for plant reproduction. In addition, our findings raise the intriguing possibility that double fertilization and seed formation may be under the influence of the circadian clock.

## Materials and Methods

### Mapping known *cis*-regulatory elements to *LORELEI* and its paralogs

Two strategies were used to identify putative CREs in *LRE* and its paralogs. In the first approach, two sets of known motif datasets were used. The first motif set includes 355 Position frequency matrices (PFMs) of Arabidopsis TFs obtained from the Cis-BP database (Weirauch et al. 2014). The PFMs were converted to position weight matrices (PWMs) with the MotifTools program in Tools for Analysis of Motifs (TAMO) (Gordon et al. 2005), which included an adjustment to the background AT (0.33) and CG (0.17) contents of the Arabidopsis genome. The second motif set was obtained from (Vandepoele et al. 2009) in the form of consensus sequences that were also converted to PWMs with TAMO. These two sets of CREs were mapped to the putative promoter (<=1kb upstream of the translation start site without including neighboring genes) and gene body sequences of *LRE*/paralog based on TAIR (http://www.arabidopsis.org) v.10 genome annotation with Motility (https://github.com/ctb/motility) and mappings with a *p*-value <1e-5 were included. If a motif did not have any mapping instances lower than this threshold, the mappings in the 90th percentile of *p*-values were included (Zou et al. 2011).

### Processing of expression datasets to identify synergid cell enriched *cis*-regulatory elements

In our second approach, we identified which CREs are enriched in synergid cell-expressed genes, by globally identifying synergid cell co-expression clusters using a compendium of four expression datasets. The first was 79 experiments in various tissues at different stages of Arabidopsis development (Schmid et al. 2005) from AtGenExpress (http://www.weigelworld.org/resources/microarray/AtGenExpress/). The second contained 36 experiments based on treatments with plant hormones (Goda et al. 2008) from the Gene Expression Omnibus (https://www.ncbi.nlm.nih.gov/geo/), accession GSE39384. The third dataset included female gametophyte cell-specific expression profiling experiments (Wuest et al. 2010) comprising the egg, central, and synergid cells from ArrayExpress (https://www.ebi.ac.uk/arrayexpress/) accessions E-MEXP-2227. Lists of the synergid cell-, the egg cell-, and the central cell-expressed genes were obtained from Table S1 from (Wuest et al. 2010). Fourth, male gametophyte-specific datasets (Borges et al. 2008), which included Arabidopsis pollen, sperm and seedling control from ArrayExpress accessions E-ATMX-35. All downloaded data in the form of CEL files were processed using the RMA function in the Affy package (Gautier et al. 2004) in R (Team). The intensity values from these four data sources were combined into an expression matrix and quantile normalized with the Affy package.

### Synergid cell co-expression clusters and TF binding motif enrichment

K-means clustering was performed using the expression matrix from the previous section with the K-means function in R iteratively such that a cluster had ≤60 but ≥10 genes. This range was determined in a previous study to balance signal-to-noise ratio and computational costs (Zou et al. 2011). Given our goal in identifying motifs controlling gene expression in synergid cells, only 5,446 genes expressed in synergid cells (Wuest et al. 2010) were included in the clustering analysis. The first round of clustering started with k=56, so the average number of genes in each cluster is ~100. Clusters with more than 60 genes were further sub-clustered with k=round ((number of genes in cluster)/(100+1)). Clusters smaller than 10 genes were not included in further analysis. This resulted in 145 non-overlapping co-expression clusters for synergid cells. For each cluster, we obtained the putative promoter sequences of genes within a cluster where the promoter was defined as 1,000 bp upstream of the transcription start site of a gene. Next, we asked which of the 355 known TFBMs (Weirauch et al. 2014) were mapped to the promoters of genes in a cluster in a significantly overrepresented manner compared to the rest of the genes in the expression matrix. For each cluster (C) and each TFBM (T) combination, a contingency table was constructed and a Fisher Exact Test was conducted (Fisher 0.1.4 package in R) to see if the number of times T was mapped to genes in C more frequently than the number of times T was mapped to genes that were not expressed in synergid cells (thus, not in any of the synergid cell co-expression clusters). A Fisher exact test was performed on each TFBM within each cluster using the Python Fisher 0.1.4 package (https://pypi.python.org/pypi/fisher/). The enrichment *p*-values were corrected for multiple testing with the *p* adjusted function in R using the Benjamini-Hochberg method (Benjamini and Hochberg, 1995). Adjusted *p*-values < 0.05 were considered significantly enriched.

### Mapping EE variants to *LRE* orthologs

*LRE orthologs from eleven species* in Brassicaceae (*Aethionema arabicum, Leavenworthia alabamica, Camelina sativa, Capsella grandiflora, Capsella rubella, Boechera stricta, Arabidopsis lyrata, Brassica rapa, Schrenkiella parvula, Sisymbrium irio, and Eutrema salsugineum*) were previously identified in (Noble et al. 2020). Promoters were considered as regions up to 1,000 bp upstream of the translation start site. EE variants were mapped to regions using Find Individual Motif Occurrences (FIMO Version 5.0.5).

### Identifying Arabidopsis *REVEILLE* genes homologous to PK02532.1, a transcription factor in *Cannabis sativa* known to directly bind to the SEEL motif *in vitro*

Querying the CIS-BP database (http://cisbp.ccbr.utoronto.ca/index.php) with the SEEL motif identified that DNA binding domain (DBD) of a transcription factor in *Cannabis sativa* (PK02532.1) that is known to bind the SEEL motif (Weirauch et al. 2014). The DBD of PK02532.1 was reported as “RESWTEPEHDKFLEALQLFDRDWKKIEAFVGSKTVIQIRSHAQKYF“ (Weirauch et al. 2014). Using this DBD of PK02532.1 as a query, we searched the CIS-BP for Arabidopsis genes containing a DBD similar to that in PK02532.1. If a DBD in other members of this MYB-related family shared ≥ 87.5% identity with the DBD in PK02532.1, it is predicted to bind to the SEE or SEEL motif (Weirauch et al. 2014). By this criterion, the CIS-BP identified Arabidopsis *REVEILLE* genes *RVE3-6* and *RVE8* as strong candidates to bind the SEE or SEEL motif. These 5 TFs are part of the 11-member RVE TF family; the remaining 6 TFs had < 87.5% homology to the DBD of PK02532.1.

### Phylogenetic relationship among *REVEILLE* genes in Arabidopsis

We obtained the protein coding sequence for PK02532.1 (Weirauch et al. 2014) and all eleven members of the RVE family protein coding sequences (https://www.arabidopsis.org/), and generated an amino acid alignment using the MUSCLE algorithm in Geneious R11.1.2 using the standard parameters (Supplementary Fig. 2). The alignment was used to build a phylogenetic tree using the RAxML plugin the Geneious R11.1.2, with the GAMMA BLOSUM62 protein model, rapid bootstrapping and search for best scoring ML tree with 100 bootstraps, starting with a completely random tree (Supplementary Fig. 1).

### Plant materials and growth conditions

Arabidopsis seeds were liquid sterilized as follows: in the following manner: 100 - 300 seeds were placed into a 1.5 mL microcentrifuge tube with 1 mL of 70 % EtOH and vortexed for three seconds at maximum speed at least three times over the course of a 3 to 5 minute period to avoid flocculating seeds from not getting sufficient exposure to the sterilizing solution. The 70 % ethanol solution was discarded and replaced with 1 mL of sterilization solution (50 % bleach, 0.2 % TWEEN-20 (Sigma-Aldrich, Catalog # P9416-100ML), then vortexed as described above. The sterilization solution was discarded, and seeds were washed three times with 1 mL of ice-cold autoclaved dH_2_O each time. Seeds were plated on ½ strength MS plates (Carolina Biological Supply Co., Catalog # 195703), with 2 % sucrose with corresponding antibiotics.

Sterilized seeds on plates were stratified for three days in the dark and at 4 °C, then plates were moved to a Percival growth chamber maintained at 21 °C with continuous light (75-100 μmol·m^-2^·s^-1^). Ten-to-fourteen-day-old seedlings were transplanted to the soil and grown in the following condition: 16 h light (100-120 μmol·m^-2^·s^-1^) at 21° C and 8 h dark at 18° C as described (Kessler et al. 2010). Columbia (Col-0) is the ecotype of all Arabidopsis seeds used in this study. *pLRE::GFP, pLREm1SEEL::GFP, pLREm2SEEL::GFP*, and *pLREΔSEEL::GFP* seeds were placed on plates containing hygromycin B (20 μg/mL; PhytoTechnology Laboratories, Catalog # H397).

The *rve* T-DNA insertion lines were obtained from the Arabidopsis Biological Resource Center: *rve1-1* (SALK_057420), *rve1-2* (SAIL_326_A01), *rve1-3* (SALK_025754C; characterized in this study for the first time), *rve5-1* (SAIL_769_A09), and *rve6-2* (SAIL_548_F12; characterized in this study for the first time). *rve1-1* seedlings were resistant to kanamycin (50 μg/mL). *rve1-2, rve5-1*, and *rve6-2* seedlings were resistant to glufosinate ammonium (10 μg/mL; Oakwood Chemical, Catalog # 044851). In *rve1-1, rve1-2*, and *rve5-1*, we confirmed the presence of at least one end of the T-DNA insertion in the respective genes (Supplementary Fig. 3A and Supplementary Fig. 4A), as was reported previously (Rawat et al. 2009; Jiang et al. 2016; Gray et al. 2017). In *rve6-2*, we sequenced both ends of the T-DNA insertion in this mutant allele (Supplementary Fig. 5A). These alleles were genotyped using the primers in Supplementary Table 4.

### Cloning transgenic constructs

The *pLREΔSEEL::GFP, pLREm1SEEL::GFP*, and *pLREm2SEEL::GFP* constructs were created by mutating the SEEL motif in the previously published *pLRE::GFP* construct (Wang et al. 2017). The single strand SEEL motif sequence (5’TAATATCT3’) in the *LRE* promoter is present in the bottom strand of the *LRE* gene unit. Mutations or deletions were introduced by PCR with PrimeSTAR® GXL DNA Polymerase (TaKaRa Bio Inc.; Catalog # R050A) using primers and DNA templates listed in Supplementary Table 4. The inserts were cloned into *pLRE::GFP* plasmid linearized with *BamHI* (NEB, Catalog # R0136) and *SalI* (NEB, Catalog # R0138) by using In-Fusion HD Cloning Plus (Clontech, Catalog # 639645), which replaced the wild-type *pLRE::GFP* transgene with the modified *pLRE::GFP* transgenes. The recombinant plasmid was transformed into Stellar™ Competent Cells (Clontech, Catalog # 636763) according to the manufacturer’s protocol and positive colonies were selected on LB plates containing kanamycin (50 μg/mL).

Transgenes in the constructs generated were sequence verified (Eton Bioscience, Inc.) before transforming into *Agrobacterium tumefaciens* (GV3101 pMP90 strain). The positive colony selected for transformation into Arabidopsis was also verified by colony PCR for the presence of the transgene.

### Plant transformation

Transformation solution containing *Agrobacterium tumefaciens* (GV3101 pMP90 strain) harboring the desired transgene was sprayed onto Arabidopsis inflorescences (Chung et al. 2000). Hygromycin-resistant T1 transformants were selected as described (Harrison et al. 2006). Briefly, T1 seeds were plated and stratified in dark at 4 °C for 2-3 days. Plates were then placed into in a Percival growth chamber set at 21 °C with continuous light (70-100 μmol·m^-2^·s^-1^) for 5-6 hours, placed in the dark, at room temperature, for 3-4 days. Plates were then returned to the Percival growth chamber and transformants were selected based on presence of true leaves, which were present only in hygromycin-resistant plants.

### Isolation of single-locus insertion lines

For each construct, at least 10 T1 hygromycin-resistant transformants were transplanted to soil. Among these lines, candidate single insertion lines were identified based on segregation of resistance to hygromycin B in T2 progeny. T1 plants are expected to be heterozygous for the transgene at the insertion locus. Therefore, T1 plants that gave rise to progeny with a 3:1 ratio of resistant to sensitive plants were considered as single-locus insertion lines. Fifteen T2 plants of candidate single-locus insertion plants were transplanted to soil and T3 selfed seeds were collected to identify homozygous lines in T3 populations. Those T2 plants that gave rise to T3 progeny that segregated 100% resistance to hygromycin B were considered to be homozygous for the transgene.

### Scoring GFP expression in mature unpollinated pistils

In order to score GFP expression in mature unpollinated pistils, we emasculated stage 12c buds (Smyth et al. 1990) and twenty-four to thirty hours after emasculation, the mature unpollinated pistils were removed from the plant and placed on a double-sided tape. Carpel walls were removed by making two incisions along the replum, using a syringe needle (27 Gauge Needle, VWR, Catalog # BD305109). The pedicel and nectaries were removed, then the transmitting tract was partially split lengthwise. Dissected samples were mounted in a 5% glycerol solution with a coverslip and GFP expression in synergid cells was scored using an epifluorescence microscope (Zeiss Axiophot) with a GFP filter (excitation HQ 470/40 and emission HQ 525/50). Pictures were acquired with Picture Frame image acquisition software (Optronics).

In each T1 line carrying *pLRE::GFP, pLRE-m1-SEEL::GFP, pLRE-m2-SEEL::GFP*, or *pLREΔSEEL::GFP* constructs, we scored GFP expression in ovules from 2-3 pistils. For data reported in Fig. 3, GFP expression in homozygous single-locus insertion T3 plants was scored in 3 plants per line and 3 pistils per plant (a total of 9 pistils). GFP expression in *rve* mutant backgrounds was scored in a similar manner. After confirming both the GFP transgene and the *rve* mutation were homozygous (by scoring resistance to hygromycin B and genotyping of the corresponding *rve* transgene, respectively) GFP expression was scored in mature unpollinated pistils. For each genotype, we scored a total of 9 pistils from 3 plants and 3 pistils per plant.

Tests of significant differences in GFP-bright expression between *pLRE::GFP* and the mutant T1 lines were determined with *Chi*-square tests using 2 x 2 contingency tables where the total number of ovules scored, and those scored GFP-bright were compared in the two genotypes.

### RNA isolation, RT-qPCR, and statistical analysis

For RT-qPCR results reported in Supplementary Figs. 3-6, we emasculated pistils and 24 hours after emasculation, in each pistil, the carpel walls, pedicel, stigma, nectaries, and style were removed and the remnants of the pistil (containing only the septum, transmitting tract, and ovules) were flash-frozen in liquid nitrogen and stored at −80 °C until RNA extraction. For each genotype, two or three biological replicates were collected and thirty mature unpollinated pistils were collected for each replicate. Since *RVE* expression is influenced by circadian rhythm, we followed specific collection times and procedures to reduce the potential influence of circadian rhythm on changes in gene expression between wild-type and mutant pistils (Rawat et al. 2009; Hsu et al. 2013). For each qPCR experiment, wild-type and mutant pistils were collected over the course of five to ten days between 10:00 AM to 2:00 PM (4 - 8 hours after plants experienced dawn in the growth chamber). Each day, a range of five to ten pistils were collected for each biological replicate, alternating the collections of biological replicates of wild-type and mutant pistils until thirty pistils were collected for each sample.

Total RNA was isolated using RNeasy Plant Mini Kit (QIAGEN, Catalog # 74904) according to manufacturer’s instructions and treated with RNase-free DNase I (Life Technologies, Catalog # AM2222) to remove residual genomic DNA Reactions were cleaned up using RNeasy MinElute Cleanup Kit (QIAGEN, Catalog # 74204) and tested for RNA integrity in Agilent Bioanalyzer 2100 (Agilent Technologies). cDNA was reverse transcribed from 1 −2.5 μg of total RNA using SuperScript™ IV First-Strand Synthesis System (ThermoFisher Scientific, Catalog # 18091050).

qPCR was performed using a Bio-Rad MyiQ2 system with a 96-well block (Bio-Rad) and SensiMix™ SYBR® & Fluorescein Kit (Bioline, Catalog # QT615-05) according to the manufacturer’s protocol. 20 μL reactions were performed with 20 ng - 60 ng of cDNA in 96-well plates (ThermoFisher Scientific, Catalog # AB1400WL) and sealed with optically clear adhesive seal sheets (ThermoFisher Scientific, Catalog # AB1170).

The following qPCR program was used for all qPCR experiments: Cycle 1: (1X) Step 1: 95.0 °C for 10:00 minutes; Cycle 2: (40X) Step 1: 95.0 °C for 10:00 minutes, Step 2: 55.0 °C for 30 seconds, Step 3: 72.0 °C for 30 seconds, Data collection and real-time analysis enabled; Cycle 3: (101X) Step 1: 45.0 °C-95.0 °C for 10 seconds, increase set point temperature after cycle 2 by 0.5 °C, Melt curve data collection and analysis enabled.

Genes of interest (*RVE1, RVE5, RVE6*, and *LRE*) and the control gene *ACTIN2/8* were amplified using primers listed in Supplementary Table 4. Ct values were normalized to *ACTIN2/8*. Relative levels of gene expression were calculated according to (Qin et al. 2009). At least two technical replicates of qPCR were performed for each qPCR experiment.

RT-qPCR expression in the mutants was represented as a fraction of that for Col-0 and all data were analyzed using these standardized values. Tests of significance of differences in expression were assessed using mixed-model analysis of variance (PROC Mixed in SAS/STAT version 9.4 software, SAS Institute Inc., 2015) with biological replicates considered a random effect and nested within mutants. Genotypes were considered fixed effects. Least squares means and model-adjusted standard errors are reported for these data, and differences among means for genotypes were compared using Tukey’s test within PROC Mixed.

### Image processing

ImageJ was used to assemble image panels, insert scale bars, and prepare Figures.

### Accession numbers

Accession numbers of the genes studied in this work are as follows: *LRE* (*At4g26466*), *RVE1* (*At5g17300*), *RVE2* (*At5g37260*), *RVE3* (*At1g01520*), *RVE4* (*At5g02840*), *RVE5* (*At4g01280*), *RVE6* (*At5g52660*), *RVE7* (*At1g18330*), *RVE7-like* (*At3g10113*), *RVE8* (*At3g09600*), *LHY* (*At1g01060*), *CCA1* (*At2g46830*).

## Acknowledgments

We thank Stacey Harmer (UC Davis) for advice on RNA isolation protocol to assess *LRE* expression in *rve* mutants and for discussions on *rve* mutants. We thank Ramin Yadegari (University of Arizona) for the epifluorescence microscope (Zeiss Axiophot) and Jeremiah Hackett (University of Arizona) for the Bio-Rad MyiQ2 qPCR system. We thank Sarah Hancock for assistance in segregation analyses and isolation of single-locus insertion lines of *pLRE::GFP* and motif deletion/alteration lines. We thank the past and present members of the Palanivelu Lab for discussions. JAN was supported by the following: IGERT Comparative Genomics Program at the University of Arizona (Award ID: 0654435); NSF Graduate Research Fellowship: Grant DGE-1143953; The University of Arizona Graduate and Professional Student Council Research and Project Grant (2015-2016); the Boynton Graduate Fellowship; and the University of Arizona Graduate College Office of Diversity and Inclusion. Additional support for this work was provided by an NSF grant to RP (IOS-1146090), as well as NSF grants (IOS-1546617, DEB-1655386, IOS-2107215) and funding from the U.S. Department of Energy Great Lakes Bioenergy Research Center (BER DE-SC0018409) to SHS.

