## Supplementary Materials for "The SEEL Motif and Members of the MYB-related REVEILLE Transcription Factor Family are Important for the Expression of *LORELEI* in the Synergid Cells of the Arabidopsis Female Gametophyte"

This Supplemental Information file consists of:

Six Supplementary Figures
Four Supplementary Tables

Submitted separately in the manuscript submission system:

Four Supplementary Dataset (excel files)

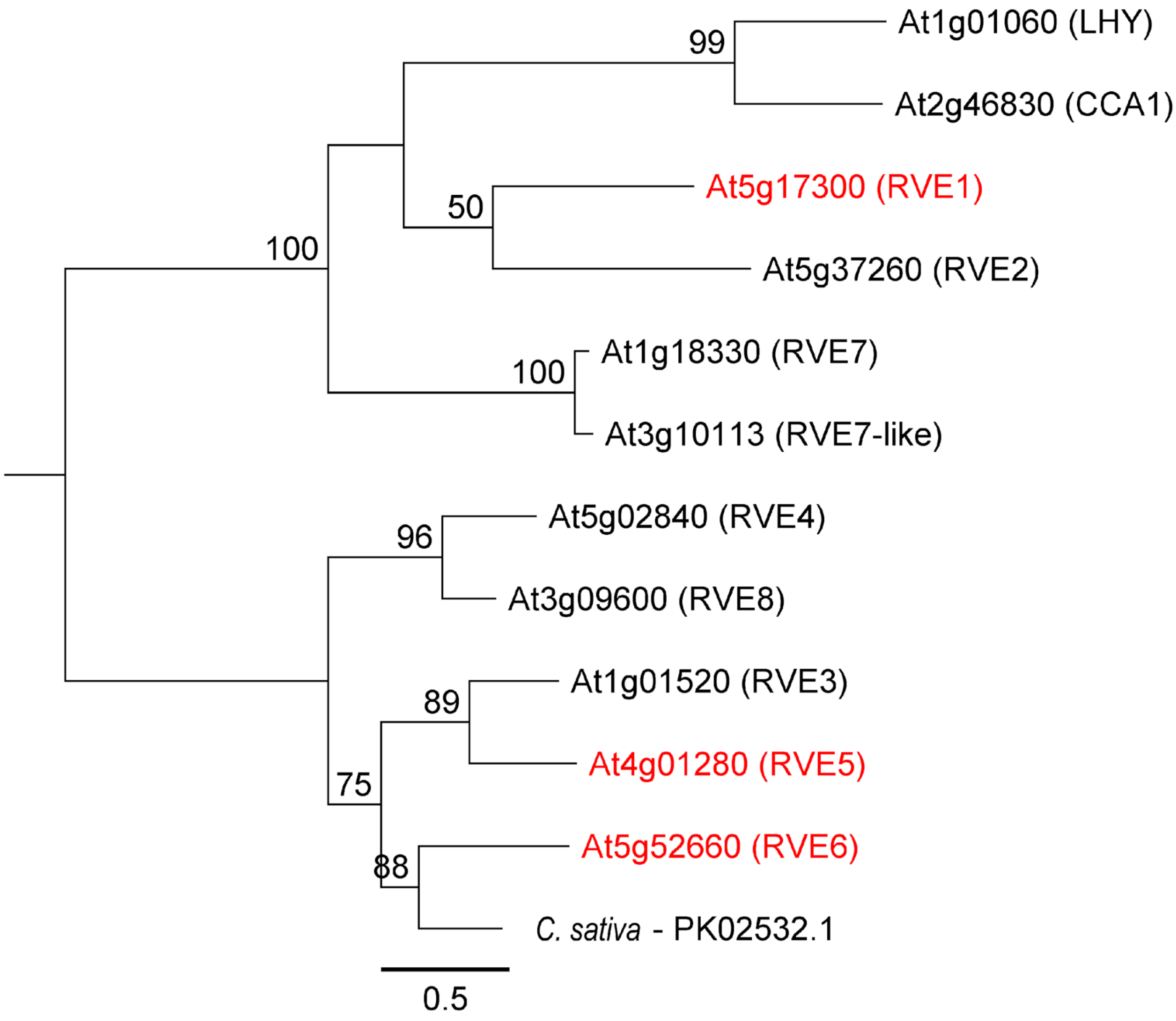


**Supplementary Fig. 1** The REVEILLE (RVE) TFs are the most closely related Arabidopsis homologs to PK02532.1, a TF in *Cannabis sativa* that directly binds to the SEEL motif *in vitro*

A phylogenetic tree of full-length amino acid sequences (Supplementary Figure 3) was built using RAxML (Geneious R11.1.2). Standard parameters were used with the GAMMA BLOSUM62 protein model, with 100 bootstrap replicates. Bootstrap values ≥ 50 are shown. Among the RVE TF family, the three RVE TFs (*RVE1*, *RVE5*, and *RVE6*) marked in red are listed as synergid-expressed in Wuest et al., 2010.

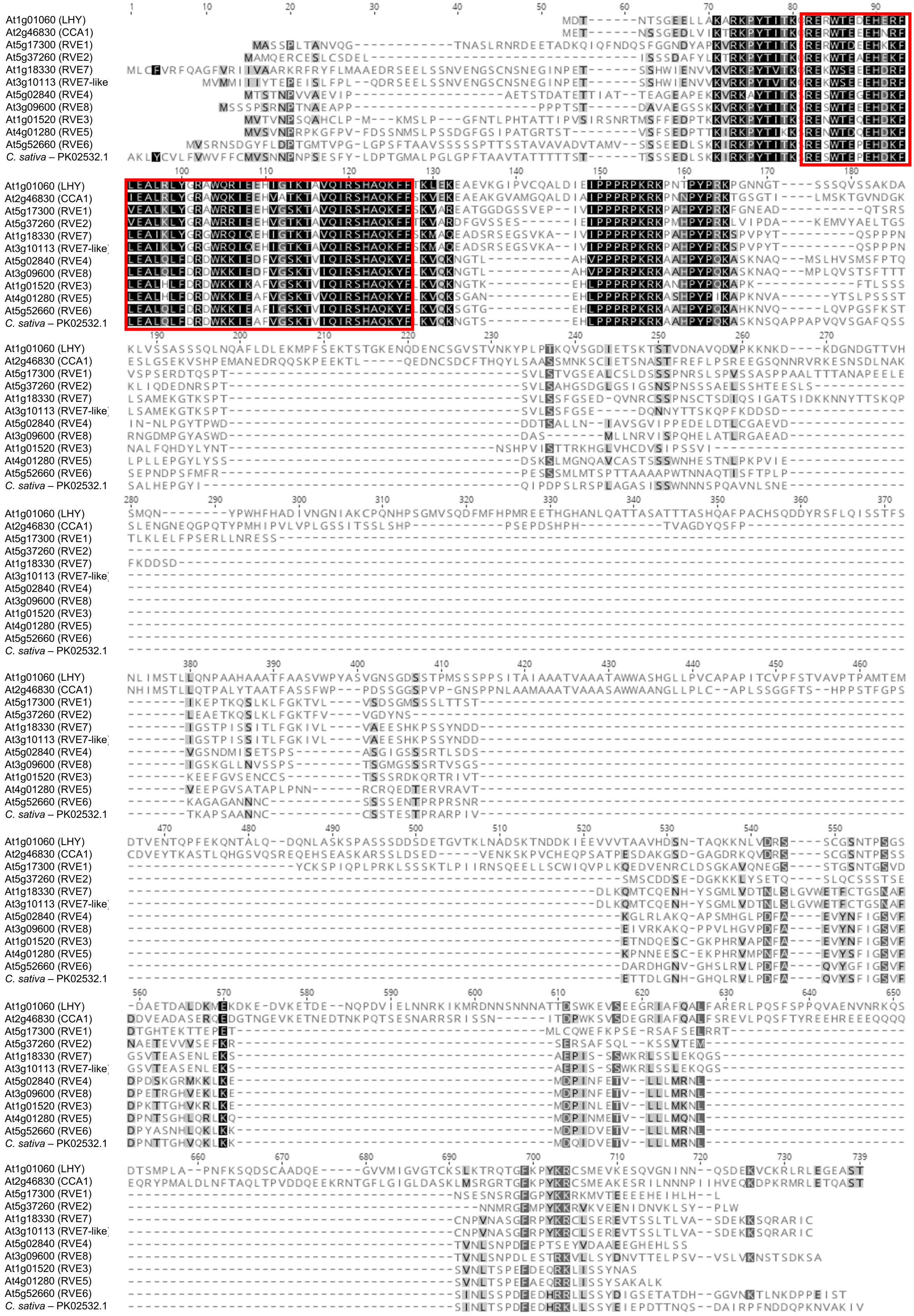


**Supplementary Fig. 2** Multiple Sequence Alignment of the REVEILLE (RVE) TF family in Arabidopsis and PK02532.1, a TF transcription factor in *Cannabis sativa* that binds to the SEEL motif *in vitro*

12 amino acid sequences were aligned using MUSCLE in Geneious R11.1.2. Amino acid sequences for PK02532.1 and 11 RVE TF family members were obtained from the CIS-BP database and TAIR, respectively. The shading gradient from black to white indicates the similarity from highest to lowest, respectively. The strongly conserved region with the SHAQKYF-type domain identified in PK02532.1 is denoted with a red rectangle.

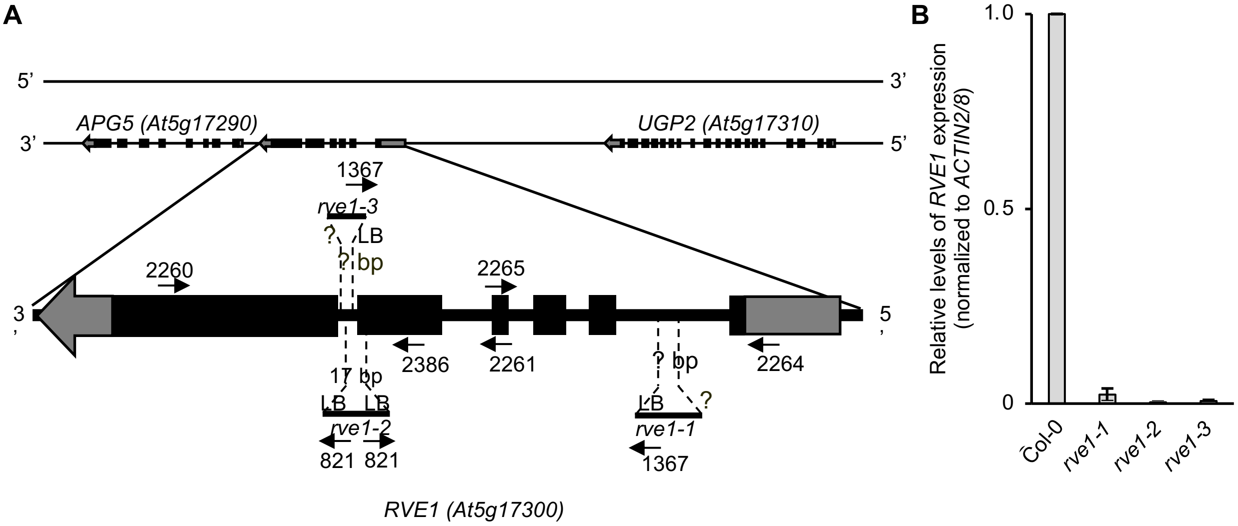


**Supplementary Fig. 3** *rve1* mutants are a knock-down alleles of *RVE1*

A. Diagram of *RVE1* gene model that also includes locations of the T-DNA insertions and primers used in genotyping, RT-PCR, and RT-qPCR. Further information about primers and PCR products can be found in Supplementary Table 4. Gray regions correspond to the 5'UTR (rectangle) and 3'UTR (arrow) of *RVE1*. The arrow points to the direction of transcription. Black rectangles refer to exons and black lines refer to introns. T-DNA insertion in *rve1-2* caused a 17bp deletion at the point of insertion; both T-DNA junctions in *RVE1* are with Left Border (LB) of the T-DNA. Only one junction between T-DNA Left Border (LB) and *RVE1* were identified in case of *rve1-1* and *rve1-3* and the extent of deletion or mutation in these insertion sites therefore remain unknown (marked with a ?).

B. RT-qPCR of *RVE1* expression in indicated mutant unpollinated pistils. Three biological replicates with two technical replicates each were used to assess *RVE1* gene expression levels. Ct values in each genotype were first normalized to *ACTIN2/8*, and then compared to wild type. Least squares means and standard errors are presented.

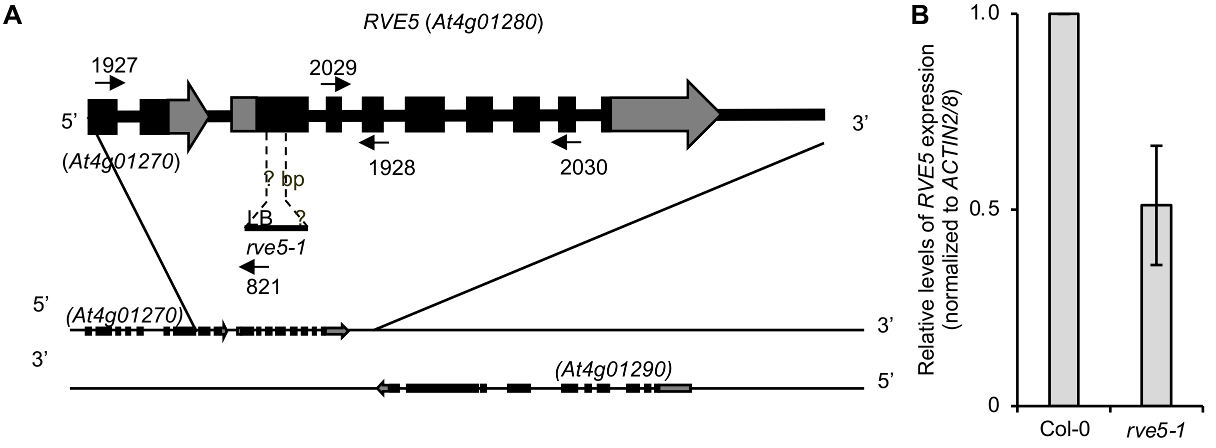


**Supplementary Fig. 4** *rve5-1* is a knock-down allele of *RVE5*

A. Diagram of *RVE5* gene model that also includes locations of the T-DNA insertions and primers used in genotyping, RT-PCR, and RT-qPCR. Further information about primers and PCR products can be found in Supplementary Table 4. Gray regions correspond to the 5'UTR (rectangle) and 3'UTR (arrow) of RVE5. The arrow points to the direction of transcription. Black rectangles refer to exons and black line refer to intron. Only one junction between T-DNA Left Border (LB) and *RVE5* was identified, hence the extent of deletion or mutation remains unknown (marked with a ?).

B. RT-qPCR of *RVE5* expression in *rve5-1* unpollinated pistils. Two biological replicates with two technical replicates each were used to assess *RVE5* gene expression levels. Ct values in each genotype were first normalized to *ACTIN2/8*, and then compared to wild type.  Least squares means and standard errors are presented.

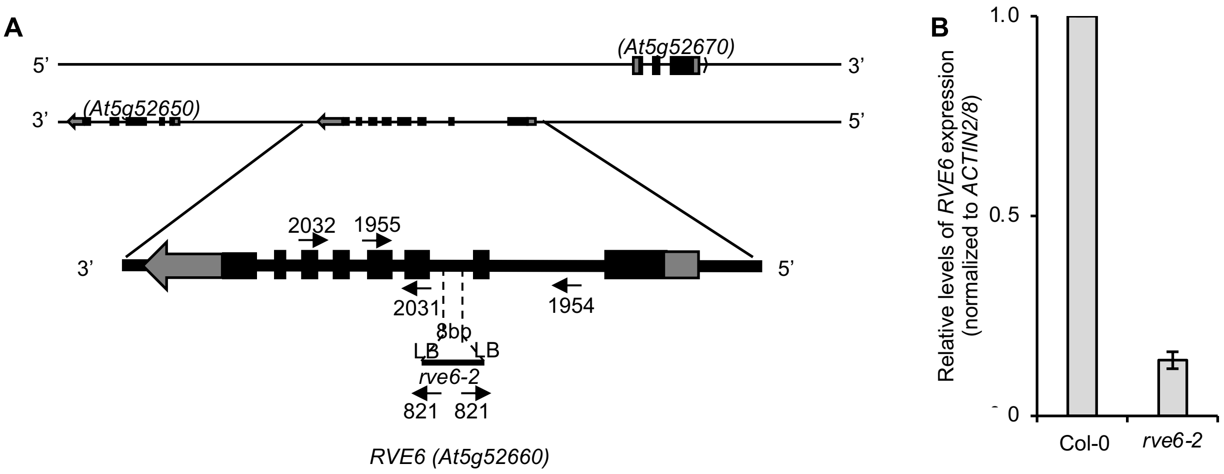


**Supplementary Fig. 5** *rve6-2* is a knock-down allele of *RVE6*

A. Diagram of *RVE6* gene model that also includes locations of the T-DNA insertions and primers used in genotyping, RT-PCR, and RT-qPCR. Further information about primers and PCR products can be found in Supplementary Table 4. Gray regions correspond to the 5’UTR (rectangle) and 3’UTR (arrow) of *RVE6*. The arrow points to the direction of transcription of *RVE6*. Black rectangles refer to exons and black line refer to intron. The T-DNA insertion in *rve6-2* caused a 8bp deletion at the point of insertion. Both T-DNA junctions with *RVE6* are with Left Border (LB) of the T-DNA.

B. RT-qPCR analysis of *RVE6* expression in *rve6-2* unpollinated pistils. Two biological replicates with two technical replicates each were used to assess *RVE6* gene expression levels. Ct values in each genotype were first normalized to *ACTIN2/8*, and then compared to wild type. Least squares means and standard errors are presented.

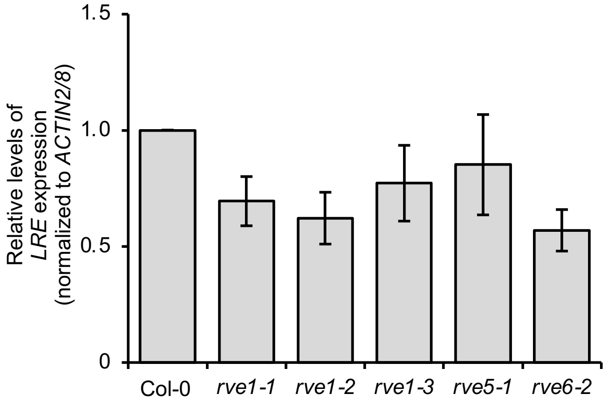


**Supplementary Fig. 6** *LRE* expression was decreased in *rve1, rve5,* and *rve6* ovules.

RT-qPCR of endogenous *LRE* expression in mature unpollinated mutant pistils compared to wild-type pistils. Ct values were first normalized to ACTIN2/8, then compared to wild-type levels. Primers spanning full length *LRE* cDNAs were used for qPCR for *rve1* mutant alleles. *LRE* expression in *rve5-1* and *rve6-2* mutant cDNAs were assessed using primers within the second and third exon of LRE. Two to three biological replicates with two technical replicates were analyzed for each mutant allele. Values were first normalized to ACTIN2/8 and then compared to wild-type levels. Least squares means and standard errors are presented.

**Supplementary Table 1** Single-locus insertion homozygous T3 lines with deletions or alterations to the SEEL motif in the promoter of the *pLRE::GFP* construct show significantly decreased percentages of GFP-positive ovules compared to that in *pLRE::GFP* lines

| Mutant genotype | Mutant line | Control line | *p*-value* | Adjusted *p*$^{+}$ |
|---|---|---|---|---|
| *pLREΔSEEL::GFP* | 22 | 1 | 1.01E-106 | 1.84E-106 |
| | 22 | 2 | 7.53E-122 | 2.82E-121 |
| | 22 | 4 | 1.44E-116 | 3.94E-116 |
| | 22 | 5 | 7.97E-109 | 1.59E-108 |
| | 22 | 10 | 3.39E-96 | 5.49E-96 |
| | 25 | 1 | 1.06E-122 | 4.89E-122 |
| | 25 | 2 | 2.43E-134 | 1.46E-132 |
| | 25 | 4 | 1.04E-128 | 1.25E-127 |
| | 25 | 5 | 2.46E-127 | 2.11E-126 |
| | 25 | 10 | 3.09E-116 | 8.06E-116 |
| | 30 | 1 | 6.08E-10 | 6.63E-10 |
| | 30 | 2 | 7.49E-14 | 9.18E-14 |
| | 30 | 4 | 2.07E-08 | 2.18E-08 |
| | 30 | 5 | 1.15E-08 | 1.23E-08 |
| | 30 | 10 | 1.95E-06 | 1.95E-06 |
| | 33 | 1 | 1.53E-117 | 4.58E-117 |
| | 33 | 2 | 3.74E-129 | 5.61E-128 |
| | 33 | 4 | 1.44E-123 | 7.85E-123 |
| | 33 | 5 | 3.42E-122 | 1.46E-121 |
| | 33 | 10 | 3.68E-111 | 7.89E-111 |
| *pLRE-m1-SEEL::GFP* | 5 | 1 | 6.36E-119 | 2.12E-118 |
| | 5 | 2 | 7.29E-131 | 1.46E-129 |
| | 5 | 4 | 2.88E-125 | 1.73E-124 |
| | 5 | 5 | 1.74E-123 | 8.69E-123 |
| | 5 | 10 | 2.84E-112 | 6.55E-112 |
| | 9 | 1 | 3.15E-13 | 3.78E-13 |
| | 9 | 2 | 2.96E-19 | 3.87E-19 |
| | 9 | 4 | 2.10E-14 | 2.63E-14 |
| | 9 | 5 | 1.02E-11 | 1.16E-11 |
| | 9 | 10 | 8.01E-08 | 8.15E-08 |
| | 12 | 1 | 4.67E-122 | 1.87E-121 |
| | 12 | 2 | 1.08E-133 | 3.24E-132 |
| | 12 | 4 | 4.55E-128 | 4.55E-127 |
| | 12 | 5 | 1.08E-126 | 7.18E-126 |
| | 12 | 10 | 1.33E-115 | 3.32E-115 |
| | 17 | 1 | 1.81E-105 | 3.20E-105 |
| | 17 | 2 | 5.41E-117 | 1.55E-116 |
| | 17 | 4 | 2.30E-111 | 5.10E-111 |

| | | | | |
|---|---|---|---|---|
| | 17 | 5 | 4.22E-110 | 8.73E-110 |
| | 17 | 10 | 3.04E-99 | 5.07E-99 |
| *pLRE-m2-SEEL::GFP* | 1 | 1 | 6.33E-52 | 8.63E-52 |
| | 1 | 2 | 1.17E-65 | 1.71E-65 |
| | 1 | 4 | 9.05E-61 | 1.29E-60 |
| | 1 | 5 | 1.33E-52 | 1.86E-52 |
| | 1 | 10 | 1.38E-41 | 1.84E-41 |
| | 7 | 1 | 1.24E-12 | 1.45E-12 |
| | 7 | 2 | 1.73E-17 | 2.21E-17 |
| | 7 | 4 | 9.57E-12 | 1.10E-11 |
| | 7 | 5 | 2.02E-11 | 2.25E-11 |
| | 7 | 10 | 2.28E-08 | 2.36E-08 |
| | 17 | 1 | 4.72E-92 | 7.26E-92 |
| | 17 | 2 | 4.54E-108 | 8.79E-108 |
| | 17 | 4 | 4.45E-103 | 7.63E-103 |
| | 17 | 5 | 7.00E-94 | 1.11E-93 |
| | 17 | 10 | 1.68E-80 | 2.53E-80 |
| | 20 | 1 | 4.89E-114 | 1.17E-113 |
| | 20 | 2 | 8.14E-127 | 6.11E-126 |
| | 20 | 4 | 2.86E-121 | 1.01E-120 |
| | 20 | 5 | 2.31E-118 | 7.31E-118 |
| | 20 | 10 | 1.01E-106 | 1.84E-106 |

Each mutant line was compared to every *pLRE::GFP* control line.
***** *p*-values were determined using a *Chi*-Square Test.
[+] *p*-values were adjusted for multiple testing with the Benjamini-Hochberg method (Benjamini and Hochberg 1995).

**Supplementary Table 2** The SEE motif is present in the promoters of other synergid cell-expressed genes that function in pollen tube reception

| Motif | Sequence Name | Start[#] | Stop[#] | Strand | Score | *p*-value | q-value | Promoter Region |
|---|---|---|---|---|---|---|---|---|
| SEE | *Arabidopsis thaliana pFER* | 717 | 724 | + | 13.43 | 5.30E-05 | 0.10 | Proximal |
| SEE | *Arabidopsis lyrata pFER* | 716 | 723 | + | 13.43 | 5.30E-05 | 0.10 | Proximal |
| SEE | *Sisymbrium irio pFER* | 758 | 765 | + | 13.43 | 5.30E-05 | 0.10 | Proximal |
| SEE | *Arabidopsis thaliana pEVN* | 490 | 497 | + | 13.43 | 5.30E-05 | 0.10 | Distal |
| SEE | *Sisymbrium irio pEVN* | 354 | 361 | - | 13.43 | 5.30E-05 | 0.10 | Distal |

[#]Start and stop refer to the nucleotide position in a promoter of a gene, which is defined as 1000bp upstream of the translation start site of that gene, except for *Sisymbrium irio pEVN* (length 910bp). Positions greater than 500 indicate presence in the proximal promoter region, beyond that they were in the distal promoter region.
Motifs were mapped using Find Individual Motif Occurrences (FIMO Version 5.0.5).
*p* values that are statistically significant and less than 1e-4 are reported.
*q* values (false discovery rate at which a motif occurrence is significant) are reported by FIMO Version 5.0.5 only if the *p* values were deemed statisically significant. They were determined by the Benjamini and Hochberg method (Benjamini and Hochberg 1995).

**Supplementary Table 3** *Chi*-square analyis of changes in *pLRE::GFP* expression in *rve* mutant ovules compared to that in wild-type ovules

| **Control Genotype** | **Mutant Genotype** | ***p*-value#** |
|---|---|---|
| GFP-bright ovules | | |
| *+/+, pLRE::GFP* | *rve1-1, pLRE:GFP* | 0.004619* |
| " | *rve1-2, pLRE:GFP* | <0.00001* |
| " | *rve5-1, pLRE:GFP* | 0.271996NS |
| " | *rve6-2, pLRE:GFP* | 0.302008NS |
| " | *rve5-1, rve6-2, pLRE:GFP* | 0.000108* |
| GFP-dim ovules | | |
| *+/+, pLRE::GFP* | *rve1-1, pLRE:GFP* | 0.000791* |
| " | *rve1-2, pLRE:GFP* | <0.00001* |
| " | *rve5-1, pLRE:GFP* | 0.27269NS |
| " | *rve6-2, pLRE:GFP* | 0.825478NS |
| " | *rve5-1, rve6-2, pLRE:GFP* | 0.008309* |
| GFP-negative ovules | | |
| *+/+, pLRE::GFP* | *rve1-1, pLRE:GFP* | <0.00001* |
| " | *rve1-2, pLRE:GFP* | <0.00001* |
| " | *rve5-1, pLRE:GFP* | <0.00001* |
| " | *rve6-2, pLRE:GFP* | <0.00001* |
| " | *rve5-1, rve6-2, pLRE:GFP* | <0.00001* |

Each mutant line was compared to a *pLRE::GFP* control line.
### *p*-values were determined using a *Chi*-Square test.
NS, *p* values Not Significant (NS) in a *Chi*-square test.
**p* values that are significant in a *Chi*-square test.

**Supplementary Table 4** List of primers used in this study

| Primer Number | Primer Description | Sequence (5' - 3') | Template | Primer Set | Expected Length (bp) |
|---|---|---|---|---|---|
| ***pLREΔSEEL::GFP plasmid*** | | | | | |
| 1935 | *pLRE::GFP* FWD Primer-1 | CGGTACCCGGGGATCCATCTGTGAGTCATCCTTTCG | *pLRE::GFP* plasmid | 1935+1936 | 747 |
| 1936 | *pLREΔSEEL::GFP* REV Primer 1 | TTGAGACCATCGTACTCTAAAACTGGCTTTG | | | |
| 1937 | *pLREΔSEEL::GFP* FWD Primer 2 | AGAGTACGATGGTCTCAAAATTTAAGGGGCCT | *pLRE::GFP* plasmid | 1937+1938 | 254 |
| 1938 | *pLRE::GFP* REV Primer 2 | TGCTCACCATGTCGACGAAATTGTTGTTAAAGAAGCTTGT | | | |
| ***pLRE-m1-SEEL::GFP plasmid*** | | | | | |
| 1935 | *pLRE::GFP* FWD Primer-1 | CGGTACCCGGGGATCCATCTGTGAGTCATCCTTTCG | *pLRE::GFP* plasmid | 1935 + 2008 | 751 |
| 2008 | *pLRE-m1-SEEL::GFP REV* Primer 1 | GACCAACTAGTCTTCGTACTCTAAAACTGGCTTTG | | | |
| 2007 | *pLRE-m1-SEEL::GFP FWD* Primer 2 | CGAAGACTAGTTGGTCTCAAAATTTAAGGGGCCT | *pLRE::GFP* plasmid | 2007 + 1938 | 256 |
| 1938 | *pLRE::GFP* REV Primer 2 | TGCTCACCATGTCGACGAAATTGTTGTTAAAGAAGCTTGT | | | |
| ***pLRE-m2-SEEL::GFP plasmid*** | | | | | |
| 1935 | *pLRE::GFP* FWD Primer-1 | CGGTACCCGGGGATCCATCTGTGAGTCATCCTTTCG | *pLRE::GFP* plasmid | 1935 + 2006 | 751 |
| 2006 | *pLRE-m2-SEEL::GFP REV* Primer 1 | GACCAGTTAAATTTCGTACTCTAAAACTGGCTTTG | | | |
| 2005 | *pLRE-m2-SEEL::GFP FWD* Primer 2 | CGAAATTTAACTGGTCTCAAAATTTAAGGGGCCT | *pLRE::GFP* plasmid | 2005 + 1938 | 256 |
| 1938 | *pLRE::GFP* REV Primer 2 | TGCTCACCATGTCGACGAAATTGTTGTTAAAGAAGCTTGT | | | |
| ***rve1* mutant genotyping** | | | | | |
| 2264 | *RVE1* genomic region left of *rve1-1* T-DNA insertion | AAGCAAATCGTTTGTTGTTGC | *Col-0* gDNA | 2264 + 2265 | 1052 |
| 2265 | *RVE1* genomic region right of *rve1-1* T-DNA insertion | GAATCTGAACTGCGGTCTTTG | | | |
| 1367 | T-DNA LB of *rve1-1* | GCGTGGACCGCTTGCTGCAACTCTCTCAGG | *rve1-1* | 1367 + 2265 | ~560 |

| | | | | | |
|---|---|---|---|---|---|
| 2261 | *RVE1* genomic region left of *rve1-2* and *rve1-3* T-DNA insertions | CAAAGACCGCAGTT CAGATTC | *Col-0* gDNA | 2261 + 2260 | 963 |
| 2260 | *RVE1* genomic region right of *rve1-2* and *rve1-3* T-DNA insertions | AACCAGTGTTTGATC CAGTCG | | | |
| 821 | T-DNA LB of *rve1-2* | GCTTCCTATTATATC TTCCCAAATTACCAA TACA | *rve1-2* | 2261 + 821; 2260 + 821 | ~750; ~650 |
| 1367 | T-DNA LB of *rve1-3* | GCGTGGACCGCTTG CTGCAACTCTCTCA GG | *rve1-3* | 2261 + 1367 | ~690 |
| ***rve5-1* mutant genotyping** | | | | | |
| 1927 | *RVE5* genomic region left of *rve5-1 T-DNA insertion* | TGAAGATTGTCCAAT GCAGGTA | *Col-0* gDNA | 1927 + 1928 | 1334 |
| 1928 | *RVE5* genomic region right of *rve5-1 T-DNA insertion* | GTCGAGGAGGTGGA AGATGT | | | |
| 821 | T-DNA LB of *rve5-1* | GCTTCCTATTATATC TTCCCAAATTACCAA TACA | *rve5-1* | 821 + 1927 | ~1100 |
| ***rve6-2* mutant genotyping** | | | | | |
| 1954 | *RVE6* genomic region left of *rve6-2 T-DNA insertion* | GTGAAGAACGAAAC AGGCAAG | *Col-0* gDNA | 1954 + 1955 | 1125 |
| 1955 | *RVE6* genomic region right of *rve6-2 T-DNA insertion* | GGAGGGGAGTGAA GCTAATTG | | | |
| 821 | T-DNA LB of *rve6-2* | GCTTCCTATTATATC TTCCCAAATTACCAA TACA | *rve6-2* | 1954 + 821; 1955 + 821 | ~850; ~550 |
| **RT-qPCR primers** | | | | | |
| 658 | *ACTIN2/8* Primer Set 1 | TCCCTCAGCACATT CCAGCAGAT | cDNA | 658 + 659 | gDNA: 155 |
| 659 | | AACGATTCCTGGAC CTGCCTCATC | | | cDNA: 69 |
| 464 | *ACTIN2/8* Primer Set 2 | CCTATTGAGCATGG TGTTGTTAGCAAC | cDNA | 464 + 465 | gDNA: 355 |
| 465 | | TGTGAGACACACCA TCACCAGA | | | cDNA: 277 |
| 2386 | *RVE1* | CACTGTTGGATCAG AAGCAT | cDNA | 2386 + 2260 | gDNA: 569 |
| 2260 | | AACCAGTGTTTGATC CAGTCG | | | cDNA: 473 |
| 2029 | *RVE5* | ATAGAAGCCTTTGTT GGATCA | cDNA | 2029 + 1928 | gDNA: 201 |
| 1928 | | GTCGAGGAGGTGGA AGATGT | | | cDNA: 112 |
| 2031 | *RVE6* | GCCGCTCATCCATA TCCTCA | cDNA | 2031 + 1955 | gDNA: 292 |
| 1955 | | GGAGGGGAGTGAA GCTAATTG | | | cDNA: 199 |

| | | | | | |
|---|---|---|---|---|---|
| 1376<br>1457 | *LRE* Primer Set 1 | ACTCACGTAAGGAC ATGCAAATTC<br>ATCACAAACCTCGG TTAGTGGAAG | cDNA | 1376 + 1457 | gDNA: 582<br>cDNA: 169 |
| 681<br>805 | *LRE* Primer Set 2 | TCAAGTCAACACTAA CAAAGCAAAAACAG CGG<br>ATGGAGCTGATATT ATTATTCTTCTTTCT GATGGC | cDNA | 681 + 805 | gDNA: 985<br>cDNA: 498 |
